# Targeted cancer ablation via programmable sensing of amplified oncogenic RNA

**DOI:** 10.64898/2026.11.02.763782

**Authors:** Meng Zhang, Yixin Hu, Sean A. Yamada-Hunter, Elena Sotillo, Haoyu Dai, Connor C. Call, Aidan M. Tousley, Stewart J. Kim, Wanting Liu, Monte M. Winslow, Crystal L. Mackall, Xiaojing J. Gao

## Abstract

Selectively eliminating cancer cells based on their oncogenic drivers holds great promise but remains challenging for targeted cancer therapy. Here, we present ORCA (Oncogenic RNA-triggered Cancer Ablation), a programmable and modular platform that couples RADAR-based detection of oncogenic RNA with conditional expression of therapeutic payloads in cancer cells. We identify RADAR sensors targeting the transcripts of endogenously amplified oncogenes, including *MYC*, *FGFR2* and wild-type *KRAS* in colorectal and gastric cancer cells. Focusing on colorectal cancer (CRC) driven by extrachromosomal DNA-amplified *MYC*, we demonstrate targeted ablation of high-*MYC* cancer cells using two orthogonal strategies: anti-CD19 CAR-T therapy and enzyme-prodrug therapy. *In vivo* delivery of ORCA using adeno-associated virus (AAV) suppresses tumor growth in a subcutaneous xenograft model and prolongs survival in a CRC liver metastasis model. ORCA establishes oncogenic RNAs as actionable inputs for targeted cancer therapy via programmable RNA sensing, thus offering a strategy to target tumors through the transcripts of otherwise difficult-to-drug oncogenes such as *MYC*.

## Introduction

A longstanding goal of cancer therapy has been to improve targeting specificity, selectively eliminating cancer cells while sparing healthy tissues (*1*). Mammalian synthetic biology has contributed toward this goal by engineering biological systems to detect cancer-specific biomarkers and activate cancer killing in response (*2*). For example, chimeric antigen receptor (CAR) T-cell therapy uses engineered T cells to recognize antigens such as CD19 on the surface of cancer cells (*3*). Synthetic biomolecular circuits that sense cell state-specific transcriptional regulation (e.g., cancer-specific promoters) or intracellular, cancer-associated biomarkers (e.g., microRNAs) have also been developed for cancer therapy (*2*, *4–6*).

These strategies, while successful at enhancing targeting specificity, face certain limitations. For instance, several of these, such as CAR-T cell therapy and microRNA-based circuits, often rely on biomarkers that are *correlative* but not *causal* in cancer development. As a result, tumor cells may evolve resistance by downregulating target expression (e.g., antigen escape) (*7*). On the other hand, identifying cancer-specific promoters or microRNA panels often requires extensive screening and validation (*8–11*), with limited generalizability across different cancer types.

To overcome these limitations, we turned to programmable RNA sensing recently developed in mammalian cells, such as RADAR (*12–14*). RADAR conditions payload translation on ADAR-mediated adenosine-to-inosine (A-to-I) editing in double-stranded RNA, enabling specific detection of endogenous RNA transcripts in mammalian cells (Figure 1a) (*12*). In the context of targeted cancer therapy, RADAR can be deployed as a highly programmable and modular “cell classifier” to not only distinguish between healthy and cancer cells based on the broadly available transcriptome data, but deliver custom therapeutic payloads selectively in cancer cells.

**Fig. 1.**
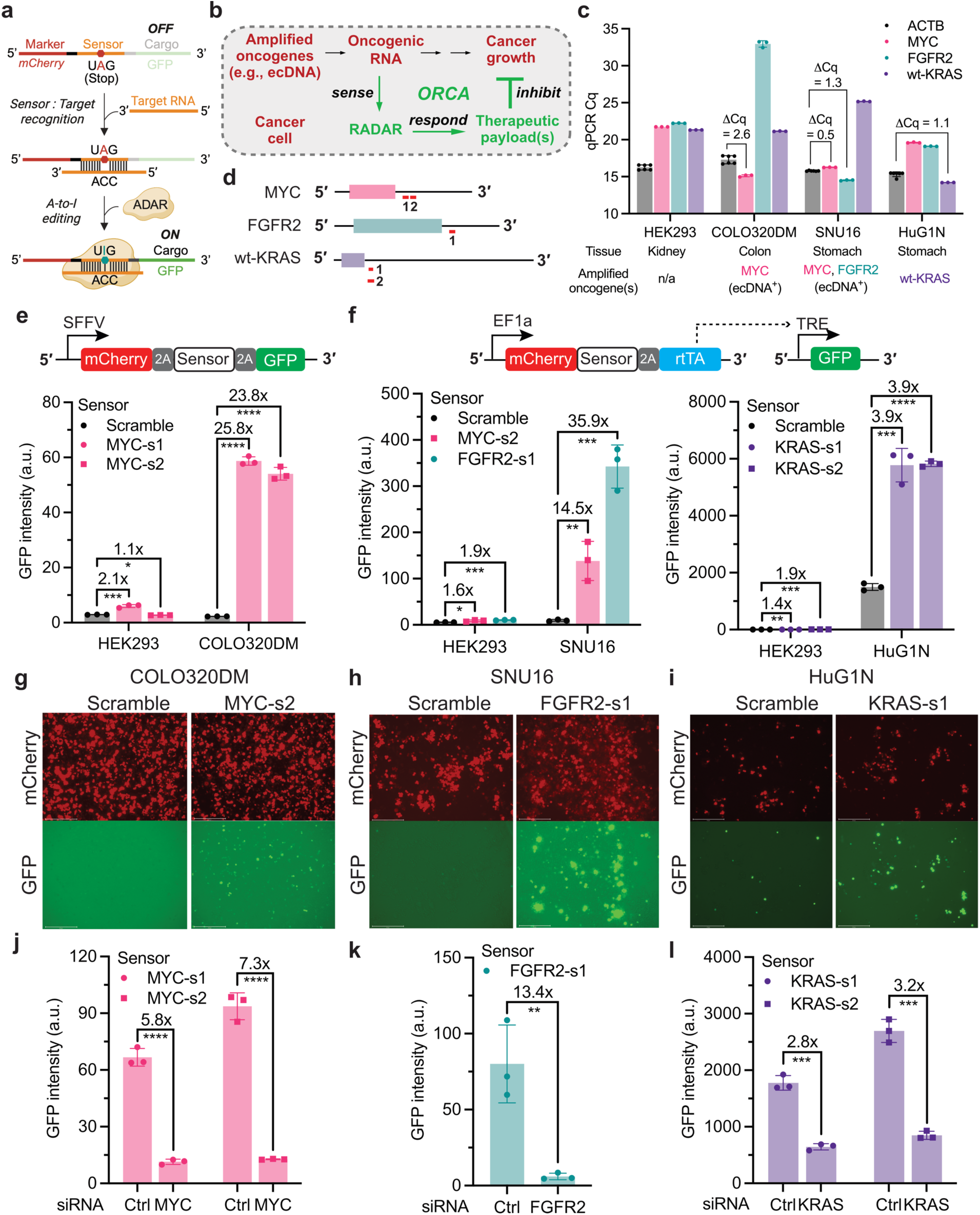
ORCA overview and characterization of RADAR sensors targeting oncogenic RNAs. **a,** RADAR mechanism. RADAR consists of three parts: an optional marker (red), a sensor sequence (orange) complementary to the target RNA except an A:C mismatch at the UAG stop codon, and a cargo (green). In the absence of target RNA, the cargo is not expressed due to the stop codon in the sensor. When sensor is bound to its target, the double-strand RNA formed between sensor and target will induce ADAR-mediated A-to-I editing, thus removing the stop codon and enabling cargo expression. **b,** ORCA overview. **c,** RT-qPCR C_q_ values of four gene targets in four cell lines. Each dot represents one technical replicate. **d**, Schematic showing targeting regions of identified sensors for each transcript. Colored box: coding sequence. Black line: 5’ or 3’ UTR. Red line: sensor target. Drawn to scale (MYC-s1 = 180 nt). **e,f**, GFP intensity of indicated cells transfected with the RADAR reporter (top) plus ADAR1p150 plasmids. Scramble sensor: a control for background payload expression. **g-i**, Representative fluorescent images of cells transfected with RADAR (shown in **e** and **f**) and ADAR1p150 plasmids. SNU16 and HuG1N are suspension cells. Images taken at one Z-section. Scale bar = 275 μm. **j-l**, GFP intensity of COLO320DM (**j**), SNU16 (**k**), and HuG1N (**l**) cells co-transfected with RADAR (shown in **e** and **f**), ADAR1p150 plasmids and siRNA. Ctrl: non-targeting siRNA. Each dot represents one biological replicate in **e**,**f**,**j**,**k**,**l**. Error bar represents standard deviation in all figures. Significance determined by two-tailed Student’s *t*-test. *: p<0.05, **: p<0.01, ***: p<0.001, ****: p<0.0001.

While RADAR and similar technologies can sense a variety of endogenous RNAs (*12–14*), we chose to focus on the transcripts of amplified oncogenes, reasoning that cancer cells are less likely to lose essential oncogenic drivers than correlative biomarkers during treatment or growth (e.g., oncogene addiction) (*15*). It has been shown that oncogene amplification often occurs on extrachromosomal DNA (ecDNA), which are found in nearly half of cancers including colorectal cancer (e.g., amplified *c-MYC*), gastric cancer (e.g., amplified *FGFR2*), and glioblastoma (e.g., amplified *EGFR*) (*16*, *17*). Among these oncogenes, *c-MYC* (or *MYC*) is of particular significance. As an important orchestrator of cancer growth (*18*), *MYC* is upregulated in up to 70% of all human cancers (*19*). While *MYC* remains difficult to drug at the protein level (*20*), its abundant mRNA transcripts in *MYC*-amplified cancer cells present a potential vulnerability that RADAR can exploit to build a sense-and-respond therapeutic pathway.

Here we present ORCA (<u>O</u>ncogenic <u>R</u>NA-triggered <u>C</u>ancer <u>A</u>blation), a RADAR-based therapeutic scheme for targeted cancer therapy (Figure 1b). We identified and characterized RADAR sensors targeting the transcripts of three endogenously amplified oncogenes (*MYC*, *FGFR2*, and wild-type *KRAS*) in colorectal and gastric cancer cell lines, including ecDNA^+^ cancer cells. As a potentially impactful test case, we focused on *MYC*-driven colorectal cancer (CRC) and demonstrated ORCA with two orthogonal therapeutic payloads, a surface protein CD19 for CAR-T therapy and an intracellular enzyme cytosine deaminase (CyD) for prodrug therapy (*21*), achieving robust and selective ablation of high-*MYC* CRC cells *in vitro*. To demarcate the current boundary of ORCA operations and guide future technology developments, we also explored its compatibility with different circuit architectures (e.g., AND logic) and delivery modalities. We further validated ORCA *in vivo* using two models, including a clinically relevant liver metastasis model as advanced-stage CRC often metastasizes to the liver and lacks effective treatment (*22*, *23*). Using adeno-associated viruses (AAVs) as the delivery vehicle, we achieved significant tumor repression in a subcutaneous tumor model and extended survival in the liver metastasis model, demonstrating the translational potential of ORCA.

## Results

### Screening and characterization of RADAR sensors targeting oncogenic RNAs

To test RADAR’s capability of sensing oncogenic RNAs (Figure 1a), we selected three representative cancer cell lines driven by endogenously amplified oncogenes, including the CRC cell line COLO320DM and the gastric cancer cell lines SNU16 and HuG1N (Figure 1c). COLO320DM and SNU16 are ecDNA^+^ cancer cells with *MYC* and *FGFR2* plus *MYC* amplification, respectively (*17*). HuG1N is a gastric cancer cell line carrying amplified wild-type (wt) *KRAS*, a chromosomal alteration found in 13% of gastric cancers with chromosomal instability (*24*). We confirmed oncogene amplification at the RNA level in these cell lines by reverse-transcription quantitative PCR (Figure 1c). Briefly, *MYC* expression was ∼6.1-fold higher than *ACTB* in COLO320DM, *FGFR2* expression was ∼2.5-fold higher than *ACTB* in SNU16, and wt-*KRAS* expression was ∼2.1-fold higher than *ACTB* in HuG1N cells (Figure 1c).

Using mCherry as a marker and green fluorescent protein (GFP) as a reporter, we screened 10-15 candidate RADAR sensors for each target and assayed sensor activity by flow cytometry. We first identified two 180-nt *MYC* sensors, MYC-s1 and MYC-s2 (Figure 1d), by testing the same reporter plasmid in high-*MYC* COLO320DM and low-*MYC* HEK293 cells (Figure 1c). Compared to the scramble sensor (a negative control for background reporter expression), MYC-s1 and MYC-s2 enabled strong GFP expression in COLO320DM cells and minimal GFP expression in HEK293 cells (Figure 1e,g). Increasing MYC-s2 to 240 nt or shortening it to 99 nt did not improve performance (Supplementary Fig. 1a). Amplicon sequencing revealed ∼5.43% and 3.89% A-to-I editing in the stop codon of MYC-s1 and MYC-s2 in COLO320DM cells transfected with RADAR reporter and ADAR1p150, respectively.

As a control for general differences between COLO320DM and HEK293 that might affect RADAR performance, we tested a previously reported sensor targeting *PCNA* (*14*), a housekeeping gene with similar expression levels in both cell lines (Supplementary Fig. 1b). The same *PCNA* sensor enabled comparable GFP expression in COLO320DM and HEK293 cells (Supplementary Fig. 1c), suggesting COLO320DM cells do not non-specifically promote RADAR activity. We further validated *MYC* sensors by knocking down MYC in COLO320DM using small interfering RNA (siRNA) (Supplementary Fig. 1d). We found a significant decrease in GFP signal from two MYC sensors following MYC knockdown (Figure 1j), demonstrating *MYC*-specific GFP expression by RADAR.

To demonstrate programmable sensing of additional oncogenic RNAs, we next identified sensors targeting endogenous *FGFR2* and wt-*KRAS* in gastric cancer cell lines SNU16 and HuG1N, respectively (Figure 1c). Because these cells grow in suspension and are more difficult to transfect than COLO320DM, we used a Tet-On-based amplification system to enhance reporter expression (Figure 1f). We identified one *FGFR2* sensor (183-nt FGFR2-s1) and two wt-*KRAS* sensors (93-nt KRAS-s1 and 183-nt KRAS-s2), each targeting the 3’ UTR of the corresponding transcript (Figure 1d). Compared to the scramble sensor, FGFR2-s1 enabled 35.9-fold higher GFP expression in high-*FGFR2* SNU16 cells and near-baseline GFP expression in low-*FGFR2* HEK293 cells (Figure 1f,h). Amplicon sequencing revealed ∼1.76% A-to-I editing in the stop codon of FGFR2-s1 in SNU16 cells stably expressing RADAR and ADAR1p150. Because SNU16 also carries MYC amplification on ecDNA (*17*), we tested MYC-s2 in this cell line. MYC-s2 enabled 14.5-fold higher GFP expression than baseline in SNU16 (Figure 1f, left), demonstrating that the MYC sensor functions across different cell types.

For wt-*KRAS*, both sensors produced 3.9-fold higher GFP expression than baseline in high-wt-*KRAS* HuG1N cells, whereas the longer sensor KRAS-s2 (183 nt) showed slightly higher GFP activation than the 93-nt KRAS-s1 in HEK293 cells (Figure 1f, right), likely due to nonspecific sensor activation. We further validated these sensors by knocking down *FGFR2* and wt-*KRAS* with siRNA in SNU16 and HuG1N cells, respectively (Supplementary Fig. 1e,f). Target knockdown significantly reduced GFP expression from each sensor (Figure 1k,l), supporting target-dependent reporter expression. Collectively, these results demonstrate RADAR-based programmable sensing of transcripts of endogenously amplified oncogenes across distinct cancer types. Notably, the five sensors identified here have relatively low GC content (36% on average; Supplementary Table 1) among the candidates screened, a feature that might have contributed to sensor performance by reducing secondary structure formation.

### ORCA enables CAR-T cell-based killing of CRC cells *in vitro*

Having identified sensors targeting specific oncogenic RNAs, we next sought to demonstrate ORCA by replacing the reporter with a therapeutic payload. We focused on *MYC*-driven CRC as the cancer model. *MYC* is overexpressed in ∼70% of CRC (*25*), particularly in metastatic CRC that lacks effective treatment (*26*). As *MYC* remains difficult to drug as a transcription factor, establishing ORCA targeting *MYC* transcripts could offer an alternative therapeutic strategy. In anticipation of delivery bottlenecks that would require ORCA-triggered death of neighboring cancer cells that do not harbor the sensor, we sought an ablation mode that engages the immune system. We chose the human surface antigen CD19 (hCD19) as our first therapeutic payload, which would enable us to repurpose FDA-approved anti-CD19 CAR-T therapy for CRC (Figure 2a).

**Fig. 2.**
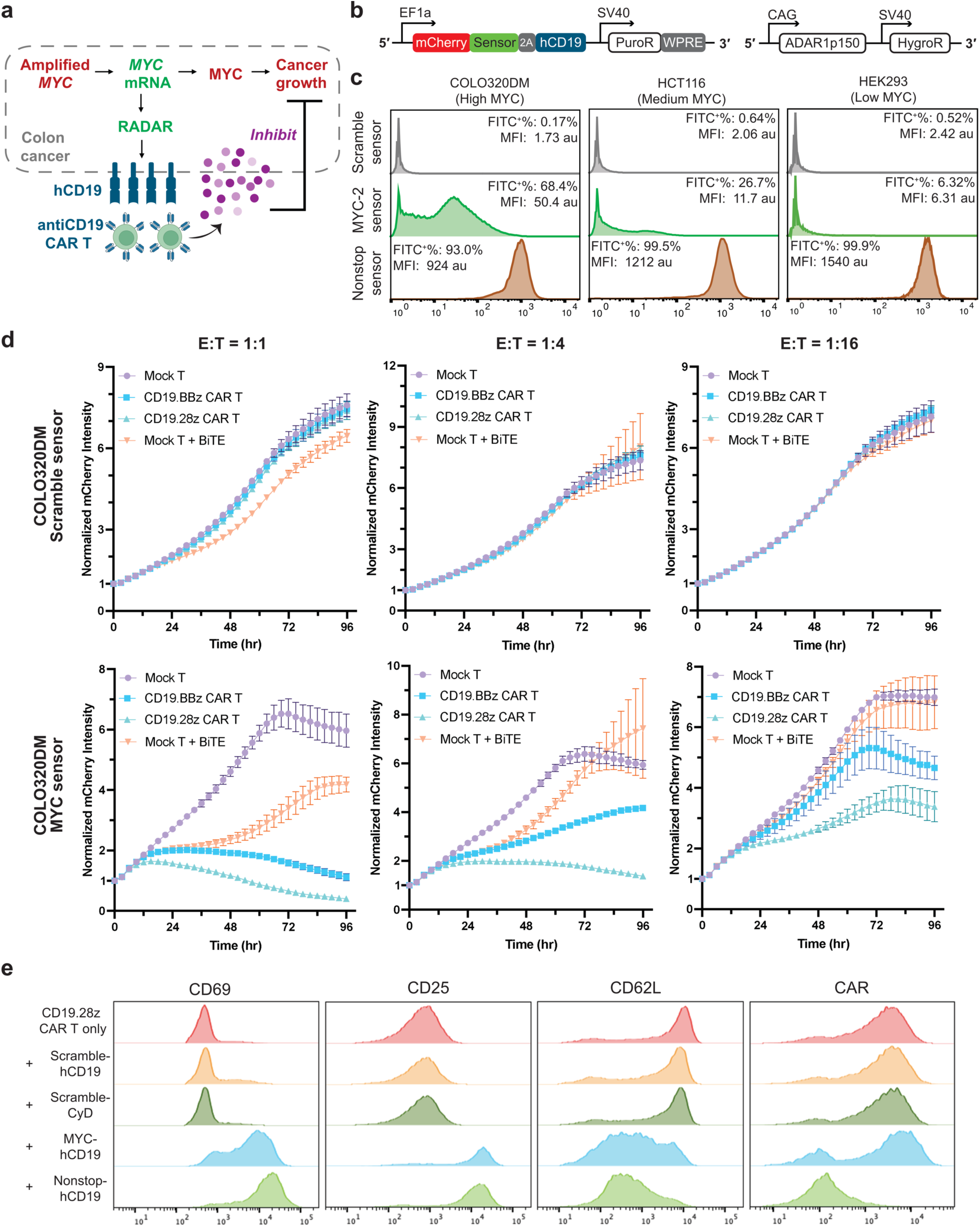
ORCA enables anti-CD19 CAR T-mediated killing of CRC cells *in vitro*. **a,** Schematic of *MYC*-specific hCD19 expression in *MYC*-amplified CRC cells for ORCA. **b**, Lentiviral vector (left) for stable expression of RADAR and PiggyBac donor vector (right) for overexpression of ADAR1p150. **c**, Histograms of hCD19 expression (determined by anti-CD19-FITC surface staining) in different cell lines stably expressing RADAR with scramble, MYC-s2, or nonstop sensor. Nonstop: control for constitutive expression where the stop codon UAG is replaced with UGG. **d**, Growth curves of engineered COLO320DM cells co-cultured with Mock T, anti-CD19-BBζ CAR-T, anti-CD19-28ζ CAR-T, or Mock T cells plus anti-CD19-anti-CD3 BiTE at varying effector-to-target (E:T) ratios. N = 3 biological replicates. Error bar represents standard deviation. **e**, Representative histograms of surface marker staining of anti-CD19-28ζ CAR-T cells at 24 hrs in the co-culture killing assay. “+ scramble-hCD19”: CAR-T cells co-cultured with engineered COLO320DM with the scramble sensor and hCD19 as payload. Similar naming applies for other conditions. CyD: cytosine deaminase, an irrelevant payload used as a control.

We selected MYC-s2 due to its lower activation in the low-MYC HEK293 cells (Figure 1e). Using transient transfection, we confirmed RADAR could deliver a surface protein hCD19 as the payload. Compared to the scramble sensor, MYC-s2 enabled significant hCD19 expression in high-*MYC* COLO320DM and minimal hCD19 expression in low-*MYC* HEK293 cells (Supplementary Fig. 2a), consistent with the GFP reporter assay. We then engineered cell lines to stably express ORCA with hCD19 as the payload and ADAR1p150 to enhance RADAR activity (Figure 2b). For comparison, we included a second CRC line HCT116 that has an intermediate level of endogenous *MYC* compared to COLO320DM and HEK293 (Supplementary Fig. 2b).

In three distinct cell lines expressing the same *MYC*-sensing ORCA at a similar rate (Supplementary Fig. 2c), we found a gradient hCD19 expression that positively correlated with their endogenous *MYC* levels. Specifically, ∼68.4%, 26.7%, and 6.32% of ORCA-engineered cells were CD19-positive in the high-*MYC* COLO320DM, medium-*MYC* HCT116, and low-*MYC* HEK293 cells, respectively (Figure 2c). In contrast, hCD19 expression under two control sensors, i.e., the *scramble* sensor for baseline expression and *nonstop* sensor for constitutive expression, was comparable across three cell types (Figure 2c). Collectively, these results suggest *MYC*-specific expression of hCD19 in multiple cell types through MYC-sensing ORCA.

To examine whether hCD19 expression level enabled by MYC-s2 is functionally adequate, we performed killing assays by co-culturing engineered COLO320DM cells with T cells and tracking cancer cell growth by Incucyte using the mCherry marker. At an 1:1 effector-to-target (E:T) ratio, we observed significant killing of *MYC*-sensing COLO320DM cells by two types of anti-CD19 CAR-T cells (BBζ and 28ζ), with CD19-28ζ outperforming CD19-BBζ CAR-T cells (Figure 2d) likely due to more potent co-stimulation (*27*). Unmodified T cells activated by an anti-CD19-anti-CD3 bi-specific T-cell engager (BiTE) also enabled significant but weaker killing of MYC-sensing COLO320DM cells. In contrast, at the same E:T ratio (1:1) we found no killing of scramble-sensing COLO320DM cells by either type of CAR-T cells, and only minor killing by BiTE-activated T cells (Figure 2d), suggesting a CD19-dependent cytotoxic response. As expected, decreasing the E:T ratio to 1:4 and 1:16 reduced the killing of *MYC*-sensing COLO320DM cells (Figure 2d), revealing a killing profile dependent on the effector-cell dose.

Surface marker staining further confirmed activation of CD19-28ζ CAR-T cells co-cultured with *MYC*-sensing COLO320DM cells, including the upregulation of CD69 and CD25, downregulation of CD62L, and a decrease in surface CAR at 24 hrs (Figure 2e). In line with surface marker activation, we found significant release of cytokines IL-2 and IFN-γ at 24 hrs by CAR-T cells (Supplementary Fig. 2d). In a tumoroid model, BiTE-activated T cells enabled significant killing of *MYC*-sensing COLO320DM cells (Supplementary Fig. 2e,f), suggesting a response in a more complex tumor microenvironment. Taken together, these results demonstrate *MYC*-sensing ORCA can drive hCD19 expression at a therapeutically meaningful level in high-*MYC* CRC cells, providing a proof of concept for CAR T-based ORCA *in vitro*.

### ORCA enables prodrug-based cancer ablation *in vitro* and *in vivo*

To expand the scope of ORCA and evaluate its compatibility with distinct therapeutic modalities, we selected cytosine deaminase (CyD), a prodrug-activating enzyme, as the second payload. CyD converts the otherwise inert prodrug 5-fluorocytosine (5-FC) into the cytotoxin 5-fluorouracil (5-FU) (*21*), a widely used chemotherapy drug particularly for CRC (*28*). We reasoned that *MYC*-sensing ORCA could enable selective CyD expression in *MYC*-amplified cancer cells, effectively turning them into “drug factories” while reducing toxicity to normal cells (Figure 3a). Additionally, the 5-FC/CyD system exhibits well-documented bystander killing effect (*29*), which could enhance its therapeutic efficacy *in vivo*.

**Fig. 3.**
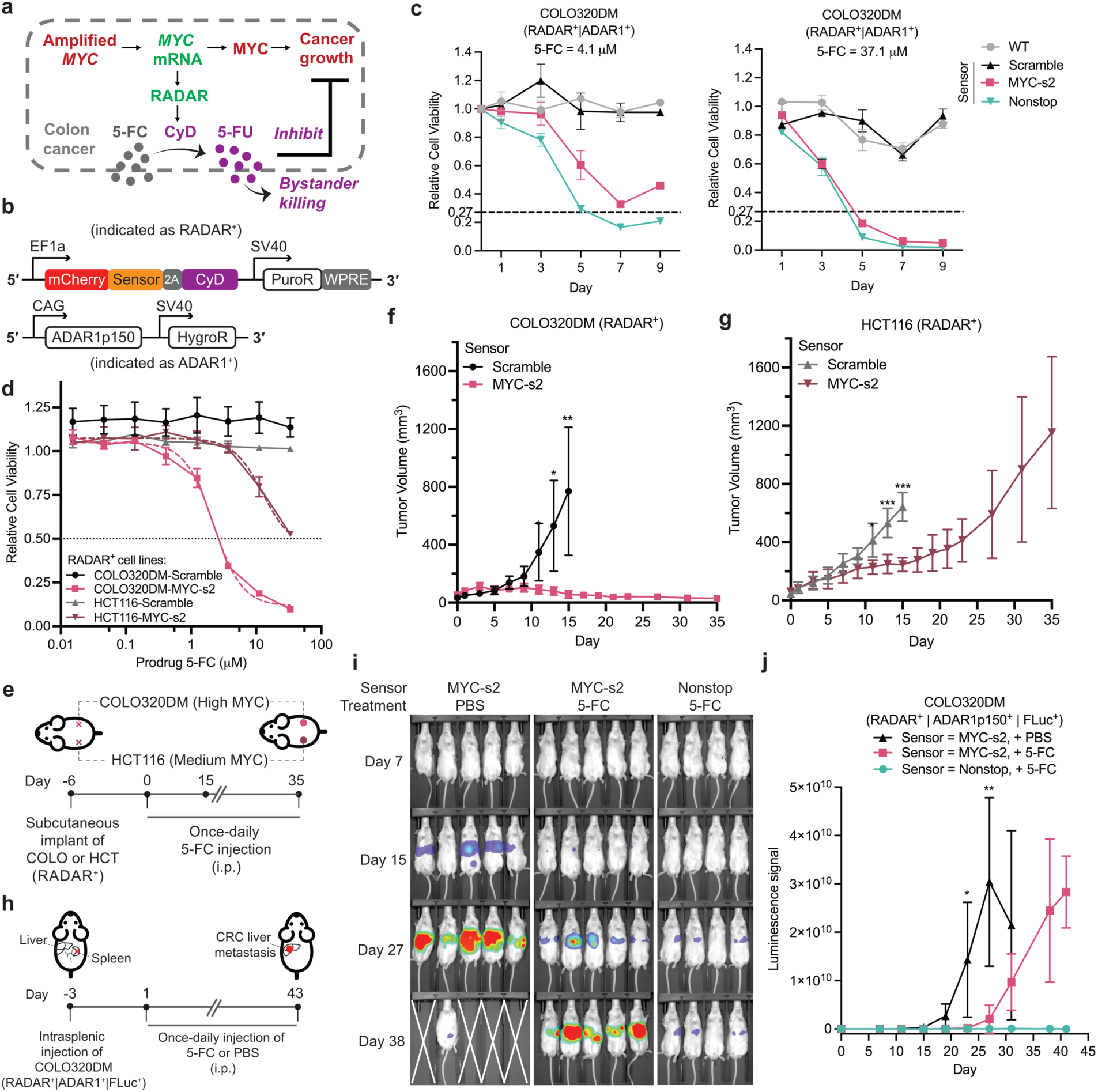
ORCA enables prodrug-based cancer ablation in vitro and in vivo. **a,** Schematic of MYC-specific expression of cytosine deaminase (CyD) for prodrug-based ORCA. **b**, Lentiviral vector (top) for stable RADAR expression and PiggyBac donor vector (bottom) for ADAR1p150 overexpression. **c,** Relative cell viability of wild-type (WT) or engineered COLO320DM cells with different sensors treated at 4.1 µM (left) or 37.1 µM (right) 5-FC. Cell viability was normalized to untreated cells on the same day. N = 3 biological replicates. **d**, 5-FC dose response curves of engineered COLO320DM or HCT116 cells expressing RADAR-CyD with MYC-s2 or scramble sensor. Cell viability was normalized to untreated cells. N = 3 biological replicates. **e**, Outline of the subcutaneous tumor model. **f,g**, Tumor growth of engineered COLO320DM (**f**) or HCT116 (**g**) cells over time. N = 5 mice. Mice carrying tumors with the scramble sensor were sacrificed on day 15 due to large tumor burden. **h**, Outline of the liver metastasis model. **i**, Bioluminescence imaging of mice in three groups on different days. **j**, Quantification of luminescence signal over time. N = 5 mice in “MYC-s2, + PBS/5-FC” groups. N = 4 mice in “Nonstop, + 5-FC” group. Error bar represents standard deviation in all figures.

We first determined the sensitivity of COLO320DM to 5-FC and 5-FU *in vitro*. Cell growth remained largely unaffected by 5-FC at concentrations up to ∼37 µM but was significantly inhibited by 5-FU, with an estimated IC_50_ of ∼10 µM (Supplementary Fig. 3a). We then generated COLO320DM cell lines stably expressing ADAR1p150 and ORCA with different sensors (Figure 3b), including MYC-s2 and two controls (scramble and nonstop). As expected, COLO320DM cells engineered with the *scramble* sensor exhibited growth curves comparable to those of wild-type cells under 5-FC treatment, whereas cells with the *nonstop* sensor showed strong growth inhibition due to constitutive CyD expression (Figure 3c). Notably, COLO320DM with MYC-s2 showed a dose-dependent response to 5-FC treatment. At a low 5-FC concentration, cell killing reached ∼67.1% by day 7, and increasing 5-FC boosted killing to ∼94.0% by day 7 (Figure 3c). Moreover, given that only ∼73% of the polyclonal COLO320DM-MYC-s2 cells were transduced with RADAR (Supplementary Fig. 3b), the observed ∼94.0% cell killing suggests a bystander killing effect.

To further demonstrate *MYC*-dependent killing by ORCA, we included HCT116 as a control for cells with intermediate endogenous *MYC* expression (Supplementary Fig. 2b). Consistent with previous reports (*30*, *31*), we found HCT116 was relatively more sensitive to 5-FU than COLO320DM, with an estimated IC_50_ of ∼5 µM (Supplementary Fig. 3c). Stable expression of *MYC*-sensing ORCA (without ADAR1p150 overexpression) in HCT116 and COLO320DM sensitized both cell lines to 5-FC treatment. Notably, at a similar transduction rate (Supplementary Fig. 3d), COLO320DM-MYC-s2 cells exhibited substantially stronger growth inhibition than HCT116-MYC-s2 cells under 5-FC treatment (Figure 3d), supporting *MYC*-dependent cell killing *in vitro*. To evaluate this effect *in vivo*, we implanted engineered COLO320DM and HCT116 cells into opposite flanks of the same immunodeficient mouse for side-by-side comparison (Figure 3e). Following once-daily intraperitoneal injection of 5-FC, we achieved robust suppression of COLO320DM-MYC-s2 tumors over 35 days (Figure 3f), whereas HCT116-MYC-s2 tumor growth was only partially inhibited (Figure 3g). As a control, COLO320DM-scramble and HCT116-scramble tumors showed comparable growth rates (Figure 3f,g). Collectively, these results demonstrate *MYC*-dependent cancer ablation *in vivo* in a subcutaneous tumor model.

While targeting a single transcript (e.g., *MYC*) enabled robust and specific cell killing by ORCA, we sought to enhance cell-type discrimination, inspired by our model CRC cell lines COLO320DM and HCT116. We explored simultaneously targeting two transcripts using AND-gate logic, a unique feature enabled by RADAR’s programmability (*12*). In addition to the quantitative difference of *MYC* levels in COLO320DM and HCT116 cells, their transcriptomic profiles also differ, offering potential targets for AND logic. Toward this goal, we identified a RADAR sensor targeting *CDX2*, a transcription factor highly upregulated in COLO320DM (*32*) but not in HCT116 cells (Supplementary Fig. 4a-d). We then designed an AND-gate sensor consisting of CDX2-s1 and MYC-s2 joined by a short 6-nt linker. Following initial screening in HEK293 cells with target overexpression (Supplementary Fig. 4e), we tested the AND-gate sensor in two CRC cell lines, using the Tet-On strategy to amplify the output. The AND-gate sensor enabled significant reporter activation in COLO320DM (6-fold higher than baseline) and near-baseline reporter expression in HCT116 cells (Supplementary Fig. 4f), demonstrating enhanced cell-type discrimination. We note, however, that the improved specificity came at the cost of decreased payload expression. Therefore, we focused on the *MYC*-sensing ORCA in following *in vivo* studies to maximize therapeutic efficacy.

We next tested the potency of prodrug-based ORCA in a CRC liver metastasis model in immunodeficient mice, a more clinically relevant system as advanced-stage CRC often metastasizes to the liver and remains difficult to treat (*22*, *23*). To evaluate performance in this challenging model without delivery restraints, we used engineered COLO320DM cells stably expressing ORCA, ADAR1p150, and firefly luciferase. Following intrasplenic injection of cells, an established method for generating CRC liver metastases *in vivo* (*33*), we initiated once-daily intraperitoneal injection of 5-FC or phosphate-buffered saline (PBS) and monitored tumor progression by bioluminescence imaging (BLI) (Figure 3h).

In the PBS-treated control group, liver BLI signals in mice began to increase 15 days after cell injection and peaked around day 27 (Figure 3i,j). These mice had a median survival of 36 days (Supplementary Fig. 5a), reflecting the aggressive nature of CRC liver metastasis. In contrast, daily 5-FC treatment significantly suppressed liver metastasis of injected COLO320DM-MYC-s2 cells for ∼27 days (Figure 3j), and all treated mice (5/5) survived through day 43 with no significant difference in body weight compared to the nonstop control group (Supplementary Fig. 5a,b). Analysis of mice confirmed liver metastases, with 5-FC treated mice showing a significantly lower liver-to-body weight ratio compared to the PBS-treated group (Supplementary Fig. 5c,d). Taken together, these results demonstrate *MYC*-sensing ORCA remained functional *in vivo*, and provide proof of concept for prodrug-based ORCA in a clinically significant CRC model. Although mice injected with COLO320DM-nonstop cells showed the lowest liver metastatic burden, we emphasize that constitutive CyD expression (i.e., nonstop sensor) lacks the cancer selectivity enabled by RADAR, and is therefore unlikely to be viable clinically.

### AAV-based delivery of ORCA suppresses tumor growth *in vivo*

To further demonstrate the translational potential of ORCA, we sought to achieve its *in vivo* delivery into established tumors using clinically approved vectors such as AAVs (*34*). We prioritized the CyD strategy, because the corresponding ORCA cassette is ∼4.3 kb in length (Supplementary Fig. 6a), which is within the 4.7 kb packaging limit of AAVs, and because it involves a simple small-molecule prodrug rather than the complex CAR-T cells. Screening of ten AAV serotypes revealed efficient transduction of COLO320DM by AAV2 and AAV1, with AAV2 outperforming AAV1 by ∼3-fold (Supplementary Fig. 6b,c). However, because AAV2 production typically yields substantially lower viral titers than other serotypes (*35*), we selected AAV1 for subsequent studies as a compromise between production practicality and transduction efficiency.

To evaluate the activity of AAV-delivered ORCA, we performed cell-killing assays in 5-FC-treated wt COLO320DM cells across a range of AAV1 doses. At a high AAV dose (10^4^ vg/cell) and an intermediate 5-FC concentration (12.3 µM), *MYC*-sensing ORCA strongly inhibited cell growth by ∼77% 5 days after transduction (Figure 4a). In comparison, the nonstop sensor performed slightly better than MYC-s2 (∼83% vs ∼77% killing), whereas the scramble sensor exhibited ∼23% cell killing (Figure 4a), likely due to background CyD expression. At the same high dose of AAV1, reducing the 5-FC concentration to 4.1 µM minimized nonspecific killing by the scramble sensor while retaining significant killing by MYC-s2 (Supplementary Fig. 6d). Notably, these experiments were performed in wt COLO320DM cells without ADAR overexpression, demonstrating that endogenous ADAR proteins are sufficient to drive significant cell killing by AAV-delivered ORCA. Moreover, a control AAV1 vector encoding *MYC*-sensing RADAR with GFP as the payload caused no significant cell growth inhibition at any dose tested (Figure 4a), indicating that RADAR-based *MYC* sensing is not inherently cytotoxic in *MYC*-amplified cancer cells.

**Fig 4.**
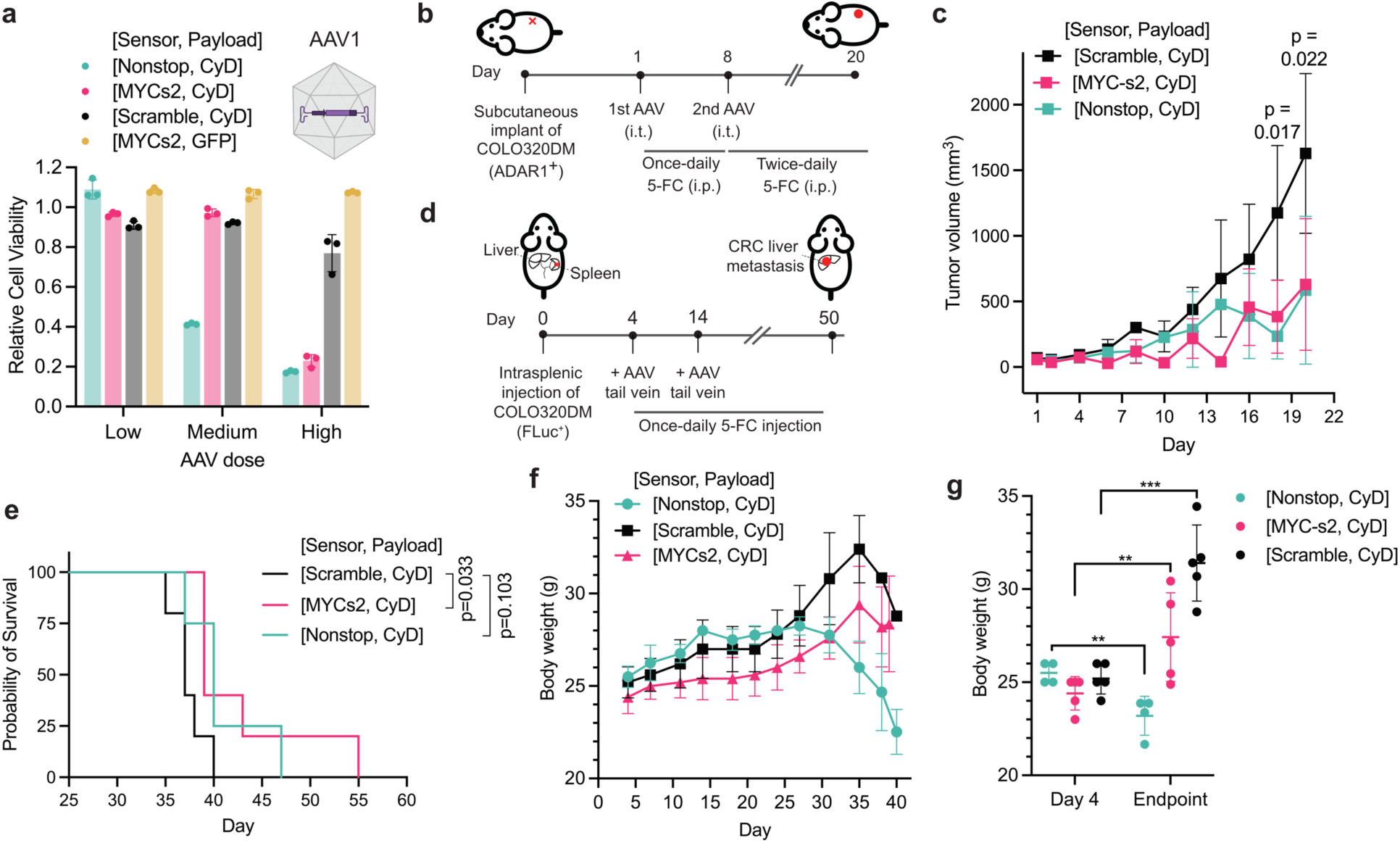
AAV delivery of ORCA suppresses tumor growth in vivo. **a**, Relative viability of wt COLO320DM cells transduced with AAV1 encoding ORCA with different sensors and payloads as indicated. Cells were treated with 12.3 µM 5-FC for 5 days following transduction. Viability was normalized to untreated cells. AAV dose: low (10^2^ vg/cell), medium (10^3^ vg/cell), high (10^4^ vg/cell). **b**, Outline of the subcutaneous tumor model with two intratumoral injections (i.t.) of AAV. ADAR1^+^ denotes cells stably expressing ADAR1p150. **c**, Subcutaneous tumor volumes over time. Day 1: first AAV injection. N = 5 mice per group (one tumor implant per mouse). P value determined by two-tailed Student’s *t*-test between the MYC-s2 group and Scramble group. **d**, Outline of the CRC liver metastasis model with two doses of AAV delivery by tail vein injection. FLuc^+^ denotes cells stably expressing firefly luciferase. **e**, Survival curves of mice in three groups. P value determined by the log-rank (Mantel-Cox) test between indicated groups. **f**, Body weight of mice over time from the first AAV injection (day 4) to day 40. **g**, Comparison of mouse body weight on day 4 and the endpoint. N = 4 in Nonstop group. N = 5 in MYC-s2 and Scramble groups. Significance determined by two-tailed Student’s *t*-test. **: p<0.01, ***: p<0.001. Error bar represents standard deviation in all figures.

A unique feature of RADAR is that it operates solely at the post-transcriptional level, making it compatible with mRNA-based delivery (*12*, *13*). In this regard, we demonstrated that *MYC*-sensing RADAR delivered as 5-methylcytosine (5mC)-modified mRNA or self-amplifying RNA (saRNA) could enable significant reporter expression in high-*MYC* COLO320DM cells (Supplementary Fig. 7a,b). However, cell killing by mRNA-delivered ORCA was lower than that achieved by AAV-based delivery, with ∼20% killing after three rounds of repeated mRNA transfection in cell culture (Supplementary Fig. 7c,d). We therefore proceeded with AAV vectors for subsequent *in vivo* studies.

In a subcutaneous tumor model using COLO320DM cells with moderate ADAR1p150 overexpression (∼2-fold above wt levels), we administered two doses of AAV1 encoding CyD-based ORCA via intratumoral injection and initiated 5-FC treatment following the first injection (Figure 4b). Compared with the scramble-sensor control, *MYC*-sensing ORCA significantly inhibited tumor growth starting on day 18 and reduced tumor volume by ∼61% by day 20 (Figure 4c). Notably, *MYC*-sensing ORCA performed comparably to the nonstop-sensor control, likely because of strong bystander killing at the relatively high prodrug concentration used *in vivo*.

We further evaluated our system in a CRC liver metastasis model using COLO320DM cells expressing firefly luciferase without ADAR1p150 overexpression. We adopted a similar two-dose regimen and administered AAV1 encoding ORCA via tail vein for systemic delivery (Figure 4d). Although mice that received *MYC*-sensing ORCA showed the lowest average BLI signals *in vivo* (Supplementary Fig. 8a-c), the differences among three groups were not statistically significant, partly due to the large variability within each group. Nevertheless, survival analysis revealed a modest but significant increase in median survival in mice treated with *MYC*-sensing ORCA compared to the scramble-sensor group (39 days vs 37 days; Figure 4e), suggesting an overall benefit. Importantly, mice receiving AAV1 with the nonstop control sensor exhibited significant body weight loss (Figure 4f,g) and signs of internal organ damage (Supplementary Fig. 8d), indicating substantial toxicity associated with constitutive CyD expression following systemic AAV delivery. In contrast, *MYC*-sensing ORCA did not cause significant body weight loss, suggesting that restricting CyD expression in high-MYC cancer cells could reduce off-target toxicity.

## Discussion

In this work, we report ORCA (Oncogenic RNA-triggered Cancer Ablation), a platform that couples programmable sensing of oncogenic RNA with conditional expression of therapeutic payloads for targeted cancer therapy. Unlike existing approaches that rely on *correlative* cancer biomarkers, ORCA achieves targeting specificity by sensing *oncogenic* transcripts essential to cancer development. Notably, ORCA is well suited against ecDNA^+^ cancers due to the abundant oncogenic RNAs present in these cells, as demonstrated here with *MYC*-amplified CRC. In this regard, ORCA also offers a strategy to address traditionally hard-to-drug oncogenes by exploiting their transcripts to establish a therapeutic pathway. Given the modular and programmable nature of RADAR, ORCA can be adapted to different cancer types by swapping the RNA sensor.

A common feature of RNA sensors identified here is their low GC content (on average ∼34% including CDX2-s1), which may reduce intramolecular secondary structure formation, thus increasing the likelihood of sensor RNA binding to its target, a critical step in sensor activation. We also demonstrated an AND-gate sensor targeting both *MYC* and *CDX2* transcripts simultaneously to discriminate two distinct CRC cell lines. Although in this work only about a dozen sensor candidates were screened for each target, high-throughput screening of a larger sensor library can provide valuable training data for deep learning language models to accelerate sensor design (*36*).

ORCA enables oncogene-dependent and tumor-localized expression of therapeutic payloads in cancer cells. We demonstrated this capability using two orthogonal, and potentially complementary, therapeutic payloads: hCD19 and CyD. While solid tumors such as CRC present several challenges for CAR T-based therapy (*37*), our strategy offers a potential solution to one key limitation, i.e., the lack of unique, clinically validated target antigens like hCD19 in solid tumors. Additionally, exploring the potential synergy between CAR-T therapy and prodrug therapy using a hCD19-CyD dual-payload system might improve ORCA efficacy, as 5-FU is known to enhance T cell-dependent antitumor immunity *in vivo* (*38*).

Efficient delivery of ORCA *in vivo* is essential for its clinical translation. ORCA delivered using AAV1 achieved significant tumor repression *in vivo*. With access to affordable large-scale production of AAV2 that transduces COLO320DM cells more efficiently than AAV1, we anticipate significant improvement in ORCA performance in CRC models *in vivo*. Alternative delivery modalities include mRNA-encapsulated lipid nanoparticles (LNPs), a clinically validated and more cost-effective platform (*39*). Systematic evaluation of chemically modified nucleosides for mRNA synthesis (*40*, *41*), as well as optimization of LNP formulations (*42*), will contribute to robust mRNA-based delivery of ORCA *in vivo*.

## Acknowledgements

This work was supported by the National Institutes of Health (NIH; R61CA278398, R21EB033858 to X.J.G.), Stanford Cancer Institute Innovation Award (X.J.G.), Stanford Canary Center Seed Grant (X.J.G.), American Cancer Society Discovery Boost Grant (X.J.G.), American Cancer Society Postdoctoral Fellowship (PF-24-1321847-01-ET to M.Z.), Parker Institute for Cancer Immunotherapy Parker Scholar Fellowship (S.A.Y-H.), CMBE Training Grant (NIH T32 GM15466301 to C.C.C), NSF-GRFP (DGE-2146755 to C.C.C.). This work was also supported by NIH grants (R35CA283888, 2P30CA124435-16 to C.L.M.), and the Virginia and D.K. Ludwig Fund for Cancer Research (C.L.M.). C.L.M. is a member of the Parker Institute for Cancer Immunotherapy, which supports the Stanford University Cancer Immunotherapy Program and is a Weill West Coast Cancer Hub Investigator. We thank Gao lab members for insightful discussion and advice. We thank Dr. Howard Chang and Dr. Paul Mischel for sharing COLO320DM and SNU16 cell lines. We thank Dr. Hyerim Yi, Dr. Jun Tang, and Dr. Shu Zhang in Mischel lab for helpful discussions. We thank Sriya Chitti (Fischbach lab) for assistance with formulating mRNA-LNPs. We thank Kevin Reynolds (Mackall lab) for assistance with CAR-T cell production. We thank Miguel Mike Alvarez and Stanford VSC for support with animal procedures, housing and husbandry. We thank Stanford Gene Vector and Virus Core (GVVC) for AAV production, and Dr. Liang Liu in Hu lab (Stanford Ophthalmology) for help with AAV packaging. We thank Dr. Nirk Quispe Calla and the team at Stanford Transgenic, Knockout and Tumor Model Center (TKTC) for assistance with intrasplenic injection. We also thank Lisa Evans at Stanford Center for Innovation in In vivo imaging (SCi³) for training and assistance with bioluminescence imaging.

## Methods

### Animals

Animal studies were conducted at Stanford University with the approval of the Stanford Administrative Panel on Laboratory Animal Care. Mice were housed in ventilated cages and subjected to a 12 h light/12 h dark cycle. Temperatures were kept between 20-22°C and relative humidity between 35-50%. Male and female immunodeficient (NSG) mice were used in the subcutaneous tumor model. Male NSG mice were used in the liver metastasis model. See detailed methods for each *in vivo* model below.

### Plasmid construction

Plasmids were generated using standard molecular cloning techniques, including In-Fusion assembly (Takara Bio, 638948) of linearized plasmids and PCR fragments, restriction–ligation of linearized fragments and annealed phosphorylated oligonucleotides. A list of plasmids used in this study are provided in Supplementary Table 3. Plasmids are available upon request and will be deposited to Addgene.

### Tissue culture

Cell lines were cultured in a humidity-controlled incubator under standard culture conditions (37 °C with 5% CO2). HEK293 cells (ATCC, CRL-1573) and COLO320DM cells (gift from Prof. Howard Chang) were cultured using DMEM (Fisher Scientific, 501015428) supplemented with 10% FBS (Fisher Scientific, FB12999102), 1 mM sodium pyruvate (EMD Millipore, TMS-005-C), 1% penicillin–streptomycin (Genesee, 25-512), and 1× MEM non-essential amino acids (Genesee, 25-536). SNU16 cells (ATCC, CRL-5974) and HuG1N cells (Creative Bioarray, CSC-C6408J) were cultured in RPMI medium (Corning, 10-040-CV) supplemented with 10% FBS and 1% penicillin–streptomycin. HCT116 cells (ATCC, CCL247) were cultured with McCoy’s 5A Medium (ATCC, 30-2007) supplemented with 10% FBS and 1% penicillin–streptomycin.

### Plasmid transfection

Adherent cells (HEK293, COLO320DM, HCT116) were seeded in 24-well tissue culture-treated plates one day before transfection. Cells were transfected the next day at around 70-90% confluency using Lipofectamine 3000 (Thermo Fisher Scientific, L3000001) with 1 µl of P3000 and 1.5 µl of Lipofectamine 3000 reagent in 50 µl of Opti-MEM (Thermo Fisher Scientific, 11058021) per 500 ng of total DNA per well in a 24-well plate. Suspension cells (SNU16 and HuG1N) were transfected using FuGENE HD (Promega, E2311) per manufacturer’s protocol. For each transfection in 24-well plate with around 150,000 cells per well, 500 ng of total DNA was added to 50 µl of Opti-MEM and mixed with 2 µl of FuGENE HD, incubated at room temperature for 5 mins, and added to each well. For the Tet-On doxycycline (dox)-inducible system, 100 ng/mL dox was added to culture media at the time of transfection.

### siRNA transfection

siRNAs targeting *c-MYC*, *FGFR2*, *KRAS*, *CDX2*, and the non-targeting control siRNA were purchased from Santa Cruz Biotechnology (catalog numbers: sc-29226, sc-29218, sc-35731, sc-43680, and sc-37007, respectively). siRNAs were resuspended at 10 µM in RNase-free water. For knockdown experiments shown in Figure 1j-l, we performed plasmid transfection first (as described above), immediately followed by siRNA transfection using Lipofectamine RNAiMAX reagent (Thermo Fisher Scientific, 13778100) per manufacturer’s protocol. Cells were analyzed ∼48 hrs after transfection.

### qPCR

Cells were trypsinized (for adherent cells), collected, and centrifuged at 400 × g for 5 mins. Total RNA was extracted using the RNeasy Mini Kit (Qiagen) with on-column DNase digestion (RNase-Free DNase Set, Qiagen) and QIAshredder (Qiagen), following the manufacturer’s instructions. RNA concentrations were measured using a NanoDrop spectrophotometer, and 2 μg of purified RNA was reverse-transcribed into cDNA using the SuperScript IV First-Strand Synthesis System (Invitrogen), with no-reverse-transcriptase controls included. Quantitative PCR was carried out on a QuantStudio 3 (Applied Biosystems) using SensiMix SYBR HiROX reagent (Meridian, QT605-05). Cq values were determined by QuantStudio Design and Analysis software. The primers used for qPCR are listed in the Supplementary Table 2.

### Flow cytometry

For transient transfections, unless otherwise described, cells were collected approximately 48 hrs after transfection, resuspended in flow buffer (culture media with 1 mM EDTA), filtered through a 40-µm strainer, and analyzed on the Bio-Rad ZE5 cell analyzer. For quantitative analysis of fluorescence intensities, data were processed using a CytoFlow Python package previously reported (*12*). When mCherry was used as a transfection marker, analyses were done by gating on mCherry-positive populations in all figures except for Supplementary Figure 4e,f where highly-transfected cells (top 5% mCherry signal) were gated for analysis. Number of replicates and statistical analysis can be found in corresponding figure legend. FlowJo software was also used for visualization and analysis of populational distributions.

### Generation of stable cell lines

PiggyBac integration was used to generate cell lines overexpressing ADAR1p150. Briefly, cells were plated in a 24-well plate and transfected the next day with 200 ng CAG-ADAR1p150-SV40-HygroR (pMZ192) and 100 ng piggyBac transposase plasmid (XG23) per well. Polyclonal cells were collected following hygromycin selection (500 μg/mL for COLO320DM, 250 μg/mL for HCT116, 200 μg/mL for HEK293) for at least 10 days. Lentiviral transduction was used to generate cell lines stably expressing ORCA with hCD19 or CyD as the payload. Lentiviral particles were produced in Lenti-X cells using a three-plasmid packaging system. Cells were seeded in 6-well plates one day before transfection and transfected at 80–90% confluency with packaging plasmids (600 ng psPAX2 and pMD2.G) and 1,100 ng of the lentiviral transfer plasmid, as described above. After 24 h, 3 mL of complete DMEM was added to each well. Viral supernatants were harvested 24 h later, passed through a 0.45-μm filter, and concentrated using Lentivirus Precipitation Solution (Alstem, VC100) according to the manufacturer’s instructions. Viral particles were collected by centrifugation at 1,500 × g for 30 mins at 4 °C and resuspended in 200 μL of complete culture medium. Concentrated lentivirus was used for transduction or stored at −80 °C until use. Approximately 1.0 – 1.5 × 10^5^ cells were seeded per well in a 24-well plate. The next day, 100-150 μL concentrated lentivirus was added per well, with an estimated MOI around 2. Polyclonal cells were collected following puromycin selection (7.5 μg/mL for COLO320DM, 1.0 μg/mL for HCT116, 0.5 μg/mL for HEK293) for at least 7 days.

### Anti-CD19 CAR-T cell production

Retrovirus was produced in 293GP packaging cells by co-transfecting anti-CD19 CAR vector plasmid with an RD114 envelope plasmid using Lipofectamine 2000. Viral supernatants were collected 48 and 72 hours after transfection, cleared by centrifugation, and stored at −80 °C. Primary human T cells were isolated from healthy donors using the RosetteSep Human T Cell Enrichment Kit with Lymphoprep density gradient medium, and cryopreserved. For CAR-T production, thawed T cells were activated on day 0 with anti-CD3/CD28 Dynabeads (1:1 bead-to-cell ratio) in RPMI medium supplemented with 10% FBS, 10 mM HEPES, 2 mM GlutaMAX, 100 U/mL penicillin and 100 µg/mL streptomycin (T cell medium) with 100 U/mL recombinant human IL-2. Non-tissue-culture-treated plates were coated overnight with RetroNectin (25 µg/mL), blocked with 2% BSA in PBS, and loaded with retroviral supernatant by centrifugation (3,200 rpm, 1.5–2 h, 32 °C). Activated T cells were added to these plates on days 2 and 3 for transduction. Beads were removed on day 4, and cells were maintained between 0.5 × 10⁶ - 2 × 10^6^ cells/mL with fresh medium every 2–3 days.

### Incucyte and data analysis

Target cells were engineered to stably express mCherry for live-cell tracking. 5 × 10⁴ target cells were seeded per well in 96-well flat-bottom plates and co-cultured in triplicate with anti-CD19 CAR-T cells (or untransduced T cells) at indicated effector-to-target ratios in 200 µL T cell medium without IL-2. Plates were imaged in an Incucyte live-cell analysis system every 3 hours for up to 96 hours, with 4 images per well at 10× magnification. Viable target cells were quantified as total integrated fluorescence intensity (or total fluorescent area, µm²/well) using the Incucyte analysis software. Values were normalized to the t = 0 measurement, and curves show the mean ± SD of triplicate wells. Graphs and statistics were generated in GraphPad Prism.

### Cytokine measurement

Cytokines IL-2 and IFN-γ were determined using Human IL-2 ELISA kit (Invitrogen, BMS221-2) and Human IFN-gamma Quantikine ELISA kit (R&D Systems, DIF50C), respectively, following manufacturer’s protocols.

### Tumoroid and BiTE killing assay

Tumoroids were generated by seeding 5,000 tumor cells per well in ultra-low attachment, round-bottom 96-well plates (Corning Costar, 7007), followed by centrifugation at 300 × g for 5 mins. After 2 days of tumoroid formation, 20,000 activated T cells and 1 ng of bispecific T-cell engager (BiTE) were added to each well, corresponding to an effector-to-target (E:T) ratio of 4:1. Tumoroids were co-cultured with T cells for 1–3 days before staining with SYTOX Green Nucleic Acid Stain (Thermo Fisher Scientific, S7020) according to the manufacturer’s instructions. Stained tumoroids were imaged, followed by flow cytometric analysis to quantify cell death.

### Prodrug killing assay

To determine the 5-FC/5-FU dose-response curves in wild-type COLO320DM and HCT116 cells, cells were plated per well at indicated seeding densities in 100 μl culture media in 96-well plates. We tested COLO320DM cells also at relatively higher seeding densities due to their smaller size compared to HCT116 cells. Culture media was changed the next day with gradient concentrations of 5-FC or 5-FU. Cells were then cultured for 3 more days. The cell viability was determined using the CellTiter-Glo 2.0 assay kit (Promega #G9241) following the manufacturer’s protocol. The relative cell viability was normalized to vehicle control.

To determine the growth curves of engineered COLO320DM cells expressing ORCA and ADAR1p150, ∼3,000 cells were plated per well in 96-well plates. The media was refreshed every other day with the indicated concentration of 5-FC, and cell viability was determined every other day using the CellTiter-Glo 2.0 assay kit (Promega #G9241). The relative cell viability was normalized to cells that received no 5-FC. To determine the 5-FC dose-response curves in engineered COLO320DM and HCT116 cells expressing ORCA, ∼3,000 cells were plated per well in 96-well plates. Culture media was changed the next day with gradient concentrations of 5-FC added. Cell viability was determined when the untreated cells reached ∼100% confluency. The relative cell viability was normalized to untreated cells.

### AAV production and screening

AAVs used in this study were purchased from Stanford Gene Vector and Virus Core (GVVC). For screening different AAV serotypes, 1 × 10^4^ wt COLO320DM cells were plated per well in 24-well plate. The next day, 3.0 × 10^9^ vg reporter AAVs (CMV-GFP) with different capsids were added per well. Cells were analyzed 6 days after adding AAV. To evaluate the killing rate of AAV-delivered circuits in wild-type COLO320DM cells, around 3,750 cells were seeded per well in 96-well plates on Day 0. AAVs were added on Day 2 at varying doses per well: 1.5 × 10^6^ vg/well (low dose), 1.5 × 10^7^ vg/well (medium dose), and 1.5 × 10^8^ vg/well (high dose). 5-FC was also added at the indicated concentrations on Day 2. The media containing 5-FC was refreshed on day 5. Cell viability was determined on Day 7 as described above. The relative cell viability was normalized to cells that received 5-FC but no AAV.

### mRNA synthesis and transfection

To synthesize linear, chemically modified mRNA: circuit-coding sequences were cloned into a plasmid backbone (gift from Dr. Michael Elowitz) to generate in vitro transcription (IVT) DNA templates by PCR. The IVT plasmid contains a T7 promoter and 5’ and 3’ UTRs optimized for expression in mammalian cells. A poly(A) tail was added to the IVT DNA template during PCR using a reverse primer containing 120 consecutive Ts. mRNA was synthesized using the CleanCap AG 3’OMe CleanScript IVT Kit (TriLink BioTechnologies, K-7413-25) and purified using the Monarch Spin RNA Cleanup Kit (NEB, T2040L) following manufacturer’s protocols. Modified nucleotide 5-Methylcytidine-5’-Triphosphate (TriLink BioTechnologies #N-1014) was used to replace CTP during IVT to reduce the immunogenicity of synthesized mRNA.

To synthesize self-amplifying RNA: template plasmids carrying the Venezuelan equine encephalitis virus (VEEV) replicon were linearized with EcoRV at 37 °C for at least 1 h and purified. In vitro transcription was performed as described above, except that CleanCap AU (TriLink BioTechnologies #N-7114) was used instead of CleanCap AG, and 5-methylcytidine-5′-triphosphate was used to replace CTP. IVT reactions were incubated at 37 °C for at least 3 h. The resulting saRNA was purified as previously described, followed by cellulose-based purification to remove double-stranded RNA (dsRNA) contaminants.

mRNA transfection was done in 96-well plate with ∼80% confluent cells using Lipofectamine MessengerMAX (Invitrogen, LMRNA008) following manufacturer’s protocol. Briefly, 100 ng mRNA was added in 10 µl of Opti-MEM and mixed with 0.3 µl of Lipofectamine MessengerMAX per well, incubated at room temperature for 5 min and added to each well. Unless otherwise noted, cells were analyzed 48 hrs after transfection.

### Firefly and NanoLuc luciferase assay

Nano-Glo Dual-Luciferase Reporter Assay system (Promega, N1610) was used to measure firefly luciferase and Nanoluc luciferase activity sequentially per manufacturer’s protocol. Luminescence was read on a TECAN Infinite M Plex microplate reader.

### CRC subcutaneous tumor model

Female and male NSG mice (6-8 weeks old) were purchased from Stanford Transgenic, Knockout and Tumor Model Center (TKTC). For side-by-side comparison (Figure 3e), engineered COLO320DM and HCT116 stably expressing RADAR (without ADAR1p150 overexpression) were used. 3 million COLO320DM cells and 2 million HCT116 cells were resuspended in 100 μL of PBS:Matrigel (1:1) and injected subcutaneously into opposite flanks of female NSG mice. I.p. injection of prodrug 5-FC was initiated 7 days after subcutaneous implant and continued daily. 5-FC dose was gradually increased from 25 mg/kg (day 1-3) to 50 mg/kg (day 4-9) to 75 mg/kg (day 10-12) and maintained at 50 mg/kg after day 12. Tumor size was measured every 2-4 days using a digital caliper. Tumor volume (V) was determined using V = 0.5 × L × W^2^.

For the subcutaneous tumor model with intratumoral AAV injections (Figure 4b), engineered COLO320DM cells moderately overexpressing ADAR1p150 (∼2-fold higher than wild-type) were used. 3 million cells were resuspended in 100 μL of PBS:Matrigel (1:1) and injected subcutaneously into the flank of male NSG mice. The first AAV i.t. injection was done when the tumor reached around 50-100 mm^3^ in size. The second AAV i.t. injection was done 14 days after subcutaneous implant. Daily 5-FC treatment (150 mg/kg, i.p. injection) was initiated 2 days after the first AAV injection and increased to twice-daily after the second AAV injection. Tumor size was measured every 2-4 days using a digital caliper. Tumor volume (V) was determined using V = 0.5 × L × W^2^.

### CRC liver metastasis model

Male NSG mice (6-8 weeks old) were purchased from Stanford TKTC. The liver metastasis model was generated at TKTC by intrasplenic injection of COLO320DM cells as previously reported (*33*). Briefly, 10^4^ cells in 50 μL culture media were injected into the mouse spleen followed by splenectomy. For the liver metastasis model with engineered cells (Figure 3h), COLO320DM cells stably expressing RADAR, ADAR1p150, and firefly luciferase were used. Once daily 5-FC or PBS treatment (i.p. injection) was initiated 4 days after surgery, with a gradual increase in dose from 25 mg/kg to 120 mg/kg. Mice were imaged by bioluminescence imaging and body weight measured every 4 days. All mice were dissected to confirm metastases at the liver. For PBS-treated mice, dissection was done at the natural endpoint when mice succumbed to tumor burden. For 5-FC-treated mice, dissection was performed on sacrificed mice on day 43.

For the liver metastasis model with systemic AAV delivery, engineered COLO320DM cells expressing only firefly luciferase were used for intrasplenic injection. The first AAV1 injection (1×10^11^ vg/mouse) was done 4 days after surgery, followed by a second AAV1 injection (4×10^11^ vg/mouse) 14 days after surgery, both by tail vein injection. Once-daily 5-FC treatment (i.p. injection) was initiated after the first AAV injection and continued throughout the study. 5-FC dose was increased from 5 mg/kg (day 4-6) to 30 mg/kg (day 7-14) to 225 mg/kg (after day 14). Mouse body weight was measured every 3-4 days and at the endpoint. Mice were imaged by bioluminescence imaging starting on day 7 and every 4-7 days afterwards. All mice were dissected at the endpoint to confirm liver metastases.

### IVIS imaging and analysis

*In vivo* bioluminescence imaging was performed using an IVIS Spectrum imaging system (PerkinElmer) with an exposure time of 2s unless otherwise noted. For firefly luciferase imaging, the substrate D-luciferin (VivoGlo Luciferin In Vivo Grade, Promega, P1041) was injected intraperitoneally at 150 mg/kg per mouse. Mice were imaged 10 min after substrate administration. Bioluminescence images were analyzed using AuraAnalysis software. Regions of interest (ROIs) of identical size were placed over the tumor-bearing regions of individual mice within each experiment, and bioluminescence signals were quantified as total photon flux (photons/s) within each ROI.

