## Supplementary Information for "Targeted cancer ablation via programmable sensing of amplified oncogenic RNA"

This document contains the following supplementary information for the submitted manuscript, including:

**Supplementary Figure 1**. RADAR sensor screening and target knockdown by siRNA.

**Supplementary Figure 2**. hCD19 as the therapeutic payload.

**Supplementary Figure 3**. Dose-response curves and RADAR marker expression in COLO320DM and HCT116 cells.

**Supplementary Figure 4**. AND-gate sensor targeting both MYC and CDX2.

**Supplementary Figure 5**. Liver metastasis model using COLO320DM cells stably expressing ORCA.

**Supplementary Figure 6**. AAV screening for ORCA delivery.

**Supplementary Figure 7**. mRNA delivery of ORCA.

**Supplementary Figure 8**. CRC Liver metastasis model with AAV-delivered ORCA.

**Supplementary Table 1**. Information of identified RADAR sensors

**Supplementary Table 2**. Primers used in this work

**Supplementary Table 3**. Plasmids used in this work

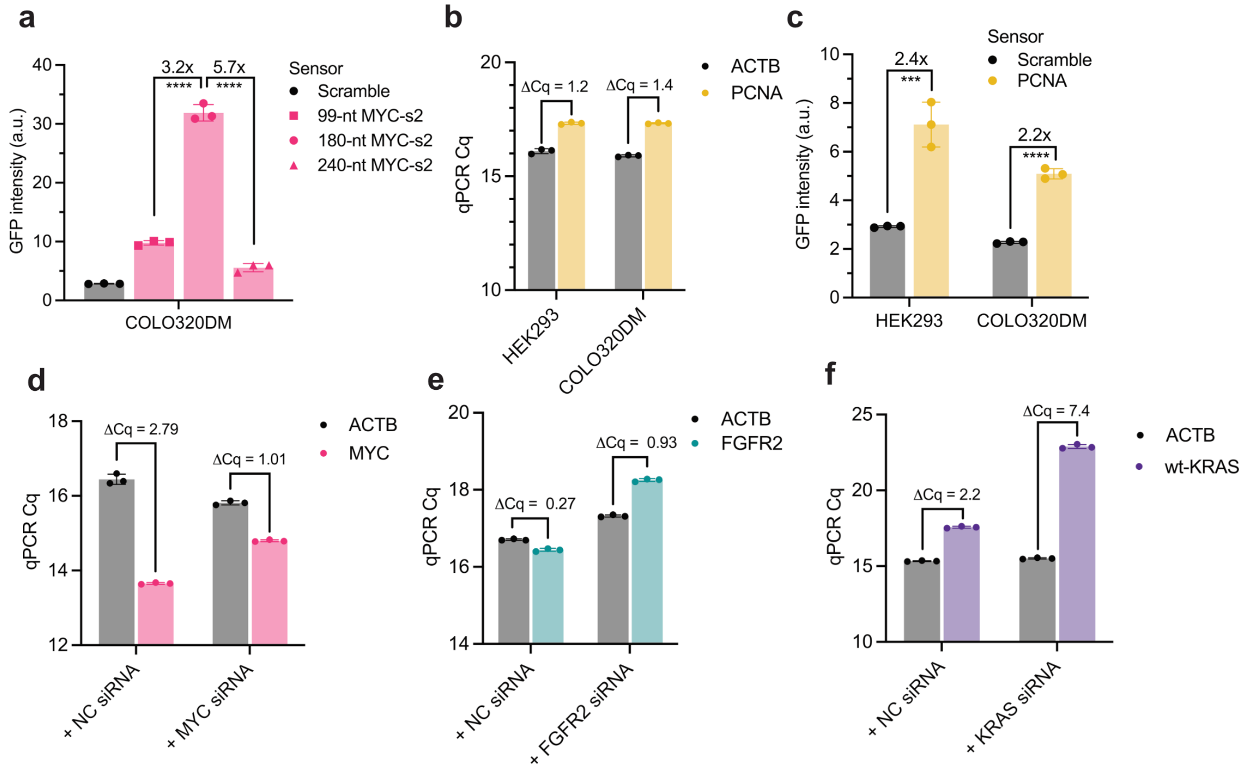


**Supplementary Figure 1. RADAR sensor screening and target knockdown by siRNA. a**, GFP intensity of cells transfected with MYC-s2 sensors of different lengths (99 nt, 180-nt, 240 nt). **b**, RT-qPCR Cq values of PCNA in COLO320DM and HEK293 cells. **c**, GFP intensity of COLO320DM and HEK293 cells transfected with the PCNA-targeting RADAR sensor. **d-f**, RT-qPCR Cq values of ACTB and target oncogenes in COLO320DM (**d**, MYC), SNU16 (**e**, FGFR2), and HuG1N (**f**, KRAS) cells treated with 5 pmol non-targeting (NC) or oncogene-targeting siRNA. Each dot represents one technical replicate. Error bars represent standard deviation in all figures. Significance determined by two-tailed Student's t-test. ***: $p<0.001$. ****: $p<0.0001$.

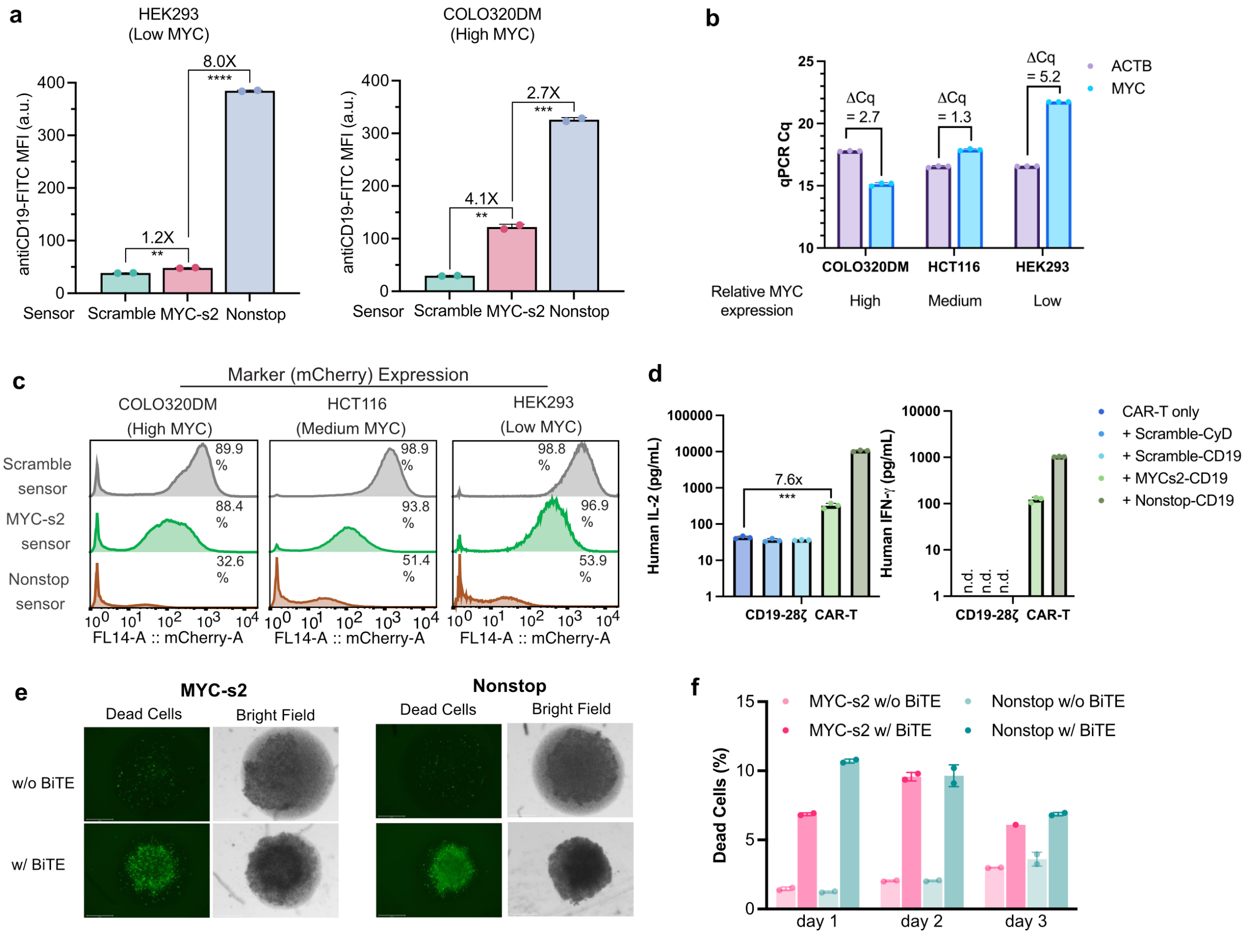


**Supplementary Figure 2. hCD19 as the therapeutic payload. a**, Anti-CD19-FITC mean fluorescence intensity (MFI) of HEK293 and COLO320DM cells transiently transfected with RADAR plasmids carrying indicated sensors and ADAR1p150 plasmid. **b**, RT-qPCR $C_q$ values of ACTB and MYC in indicated cell lines. **c**, Representative histograms of mCherry expression in cells stably expressing RADAR with different sensors and ADAR1p150 overexpression. **d**, IL-2 and IFN-γ secretion by anti-CD19-28ζ CAR-T cells after 24 hrs of co-culture with engineered COLO320DM cells. n.d.: not detected (below detection limit) **e**, Representative fluorescence and bright-field images of COLO320DM tumoroids co-cultured with T cells with or without BiTE for 24 hrs. Scale bar = 650 μm. **f**, Percentage of GFP-positive (i.e. dead) cells in tumoroids over three days of co-culture. Each dot represents one biological replicate. Error bars represent standard deviation in all figures. Significance was determined by two-tailed Student's t-test. **: p<0.01, ***: p<0.001, ****: p<0.0001.

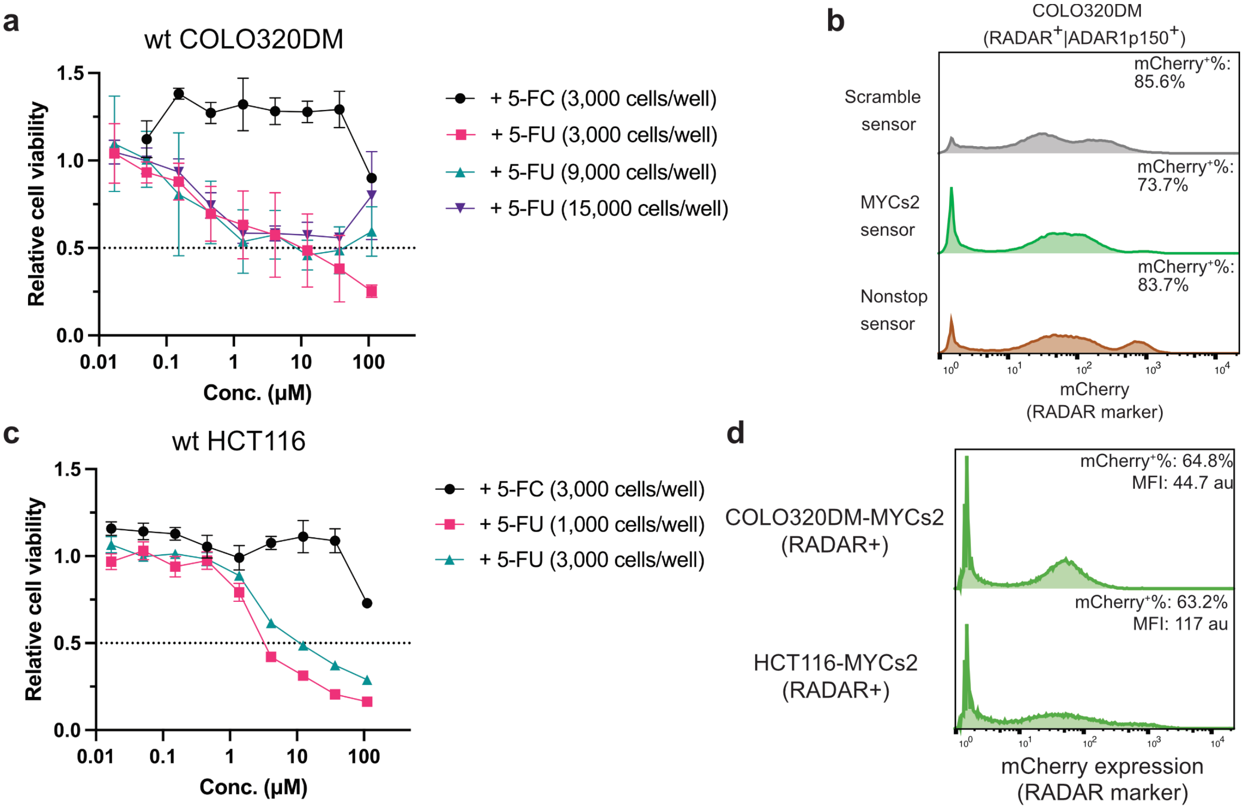


**Supplementary Figure 3. Dose-response curves and RADAR marker expression in COLO320DM and HCT116 cells.** **a**, Dose-response curves of wild-type COLO320DM cells treated with different concentrations of 5-FC or 5-FU at the indicated cell seeding densities. Cell viability was normalized to untreated cells. **b**, Representative histograms of mCherry (RADAR marker) expression in COLO320DM cells stably expressing ORCA (CyD payload) with scramble, MYC-s2, or nonstop sensor and ADAR1p150. Percentages indicate mCherry-positive cells. **c**, Dose-response curves of wild-type HCT116 cells treated with different concentrations of 5-FC or 5-FU at the indicated cell seeding densities. Cell viability was normalized to untreated cells. **d**, Representative histograms of mCherry (RADAR marker) expression in COLO320DM and HCT116 cells stably expressing ORCA (CyD payload) with the MYC-s2 sensor without ADAR1p150 overexpression. Percentages indicate mCherry-positive cells. MFI: mean fluorescence intensity. N = 3 replicates in **a** and **c**. Error bars represent standard deviation in all figures.

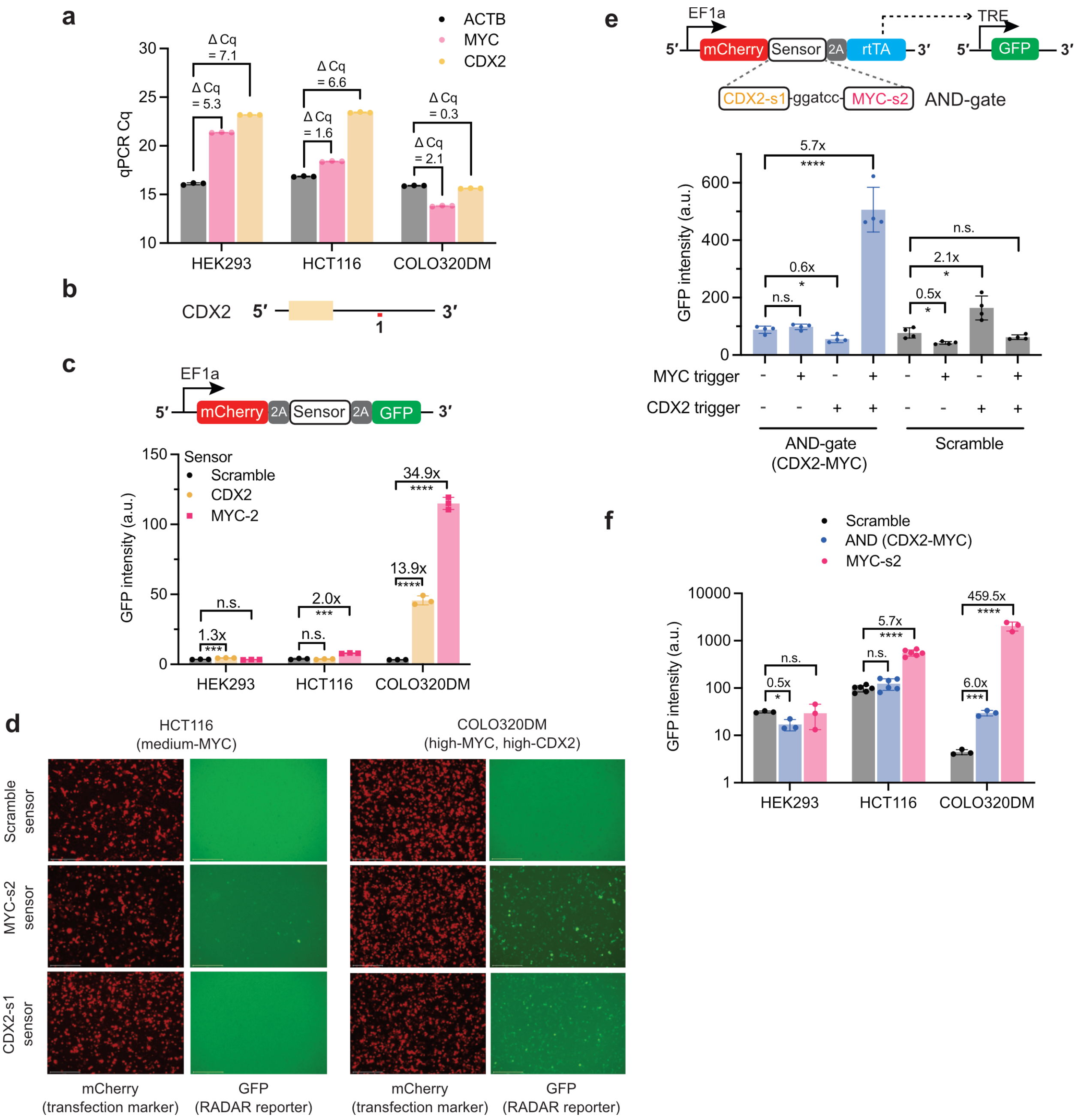


**Supplementary Figure 4. AND-gate sensor targeting both MYC and CDX2. a**, RT-qPCR $C_q$ values of ACTB, MYC, and CDX2 in indicated cell lines. **b**, Schematic showing the targeting region of CDX2-s1 on the CDX2 transcript. **c**, Schematic of RADAR reporter (top) and GFP intensity of indicated cells transfected with RADAR plasmids carrying different sensors (bottom) and ADAR1p150 plasmid. **d**, Representative fluorescence images of cells transfected with RADAR plasmids shown in **c**. Scale bar = 275 μm. **e**, Schematic of RADAR reporter with CDX2-MYC AND-gate sensor (top) and GFP intensity of HEK293 cells co-transfected with indicated sensors and triggers (bottom). **f**, GFP intensity of indicated cells transfected with RADAR plasmids carrying different sensors and ADAR1p150 plasmid. Each dot represents one replicate. Error bars represent standard deviation in all figures. Significance was determined by two-tailed Student's t-test. n.s.: not significant. *: p<0.05, ***: p<0.001, ****: p<0.0001.

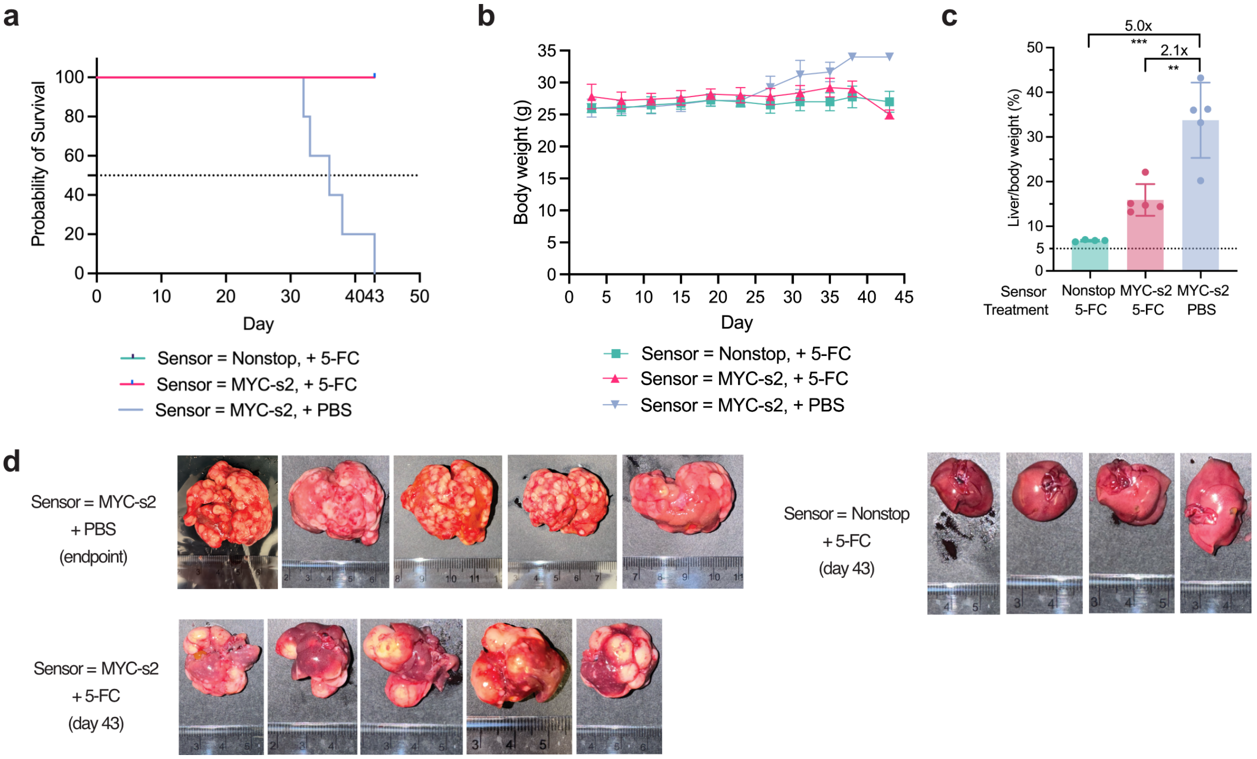


**Supplementary Figure 5. Liver metastasis model using COLO320DM cells stably expressing ORCA.** **a**, Survival curves of mice in the CRC liver metastasis model with indicated sensors and treatments. **b**, Body weight of mice over time. **c**, Liver-to-body weight ratios of mice in the indicated groups. Liver was isolated at the endpoint for each mouse in the PBS-treated group. For 5-FC-treated groups, livers were isolated on day 43 after sacrifice. **d**, Images of livers from mice in different groups. N = 4 mice in the Nonstop group and N = 5 mice in the MYC-s2 groups. Error bars represent standard deviation in all figures. Significance in **c** was determined by two-tailed Student's *t*-test. **: p<0.01, ***: p<0.001.

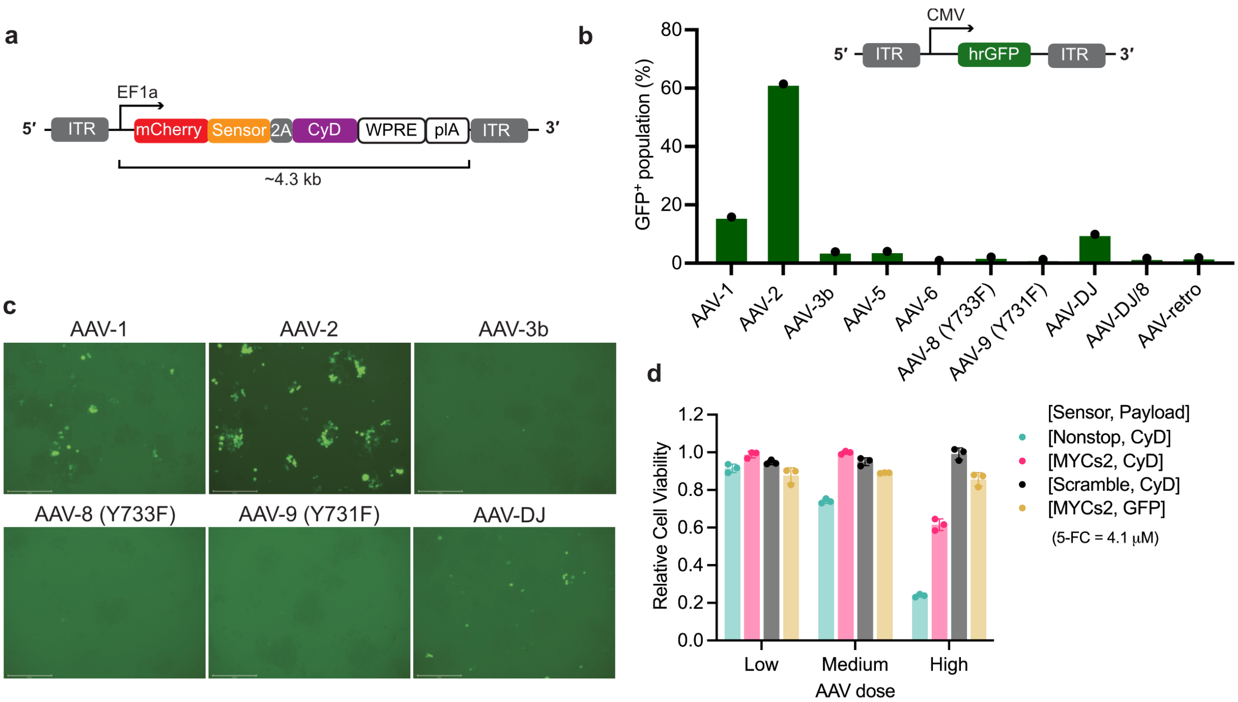


**Supplementary Figure 6. AAV screening for ORCA delivery.** a, Schematic of the AAV vector encoding ORCA with CyD as payload (~4.3 kb). **b**, Percentage of GFP-positive COLO320DM cells transduced with different AAV serotypes encoding CMV-GFP. **c**, Representative fluorescence images of COLO320DM cells transduced with indicated AAV serotypes. Scale bar = 275 μm. **d**, Relative viability of wild-type COLO320DM cells transduced with AAV1 encoding ORCA with different sensors and payloads. Cells were treated with 4.1 μM 5-FC for 5 days following transduction. Viability was normalized to untreated cells. AAV dose: low ($10^2$ vg/cell), medium ($10^3$ vg/cell), high ($10^4$ vg/cell). Each dot represents one replicate. Error bars represent standard deviation.

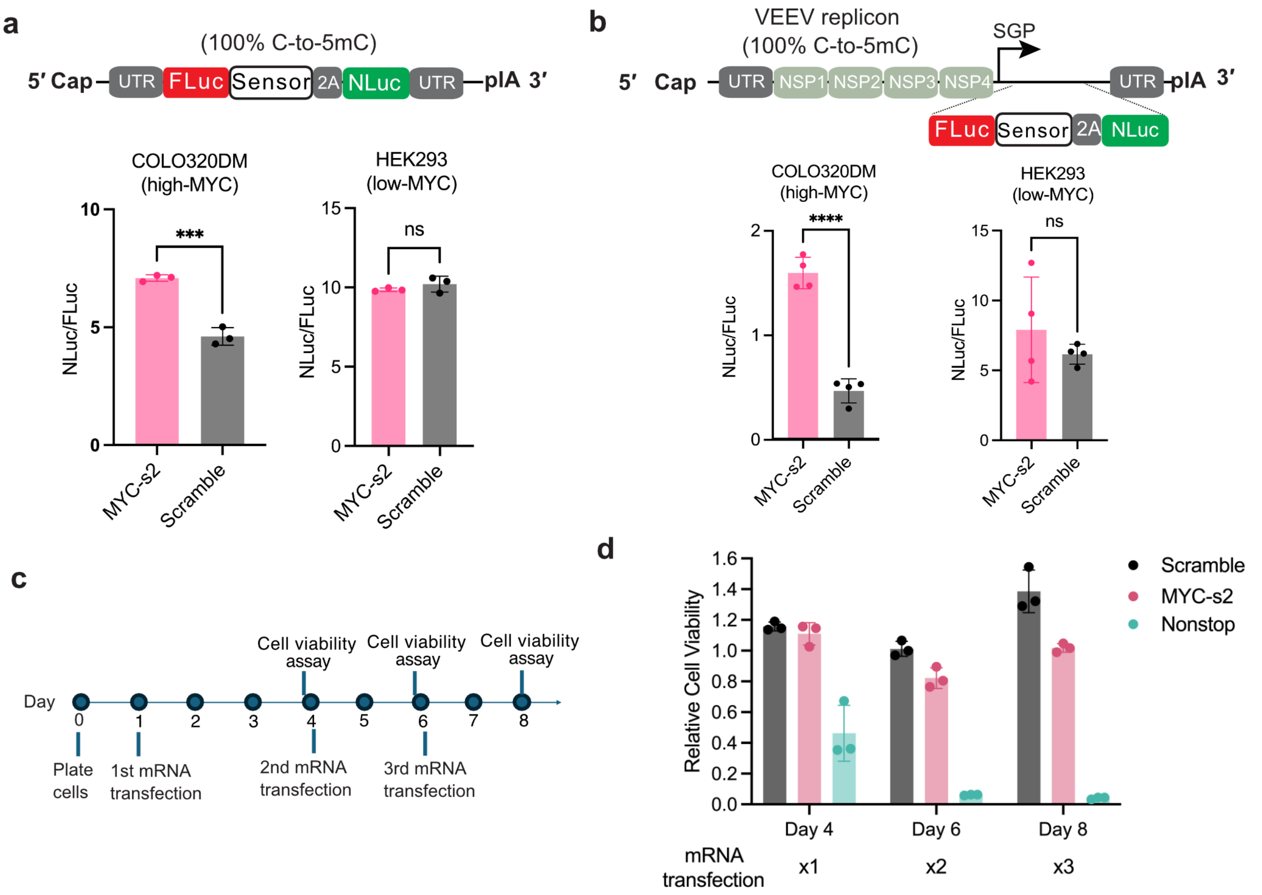


**Supplementary Figure 7. mRNA delivery of ORCA.** **a**, **b**, Schematic of 5-methylcytosine (5mC)-modified mRNA (**a**) and self-amplifying RNA (saRNA, **b**) encoding RADAR with firefly luciferase (FLuc) as transfection marker and NanoLuc luciferase (NLuc) as reporter (top). NLuc/FLuc ratios of COLO320DM and HEK293 cells transfected with indicated RADAR sensors (bottom). **c**, Timeline of repeated mRNA transfection and cell viability assays. **d**, Relative viability of COLO320DM cells transfected with mRNA encoding ORCA (CyD as payload) with different sensors following one, two, or three rounds of transfection. Each dot represents one replicate. Error bars represent standard deviation in all figures. Significance was determined by two-tailed Student's *t*-test. n.s.: not significant. ***: $p<0.001$, ****: $p<0.0001$.

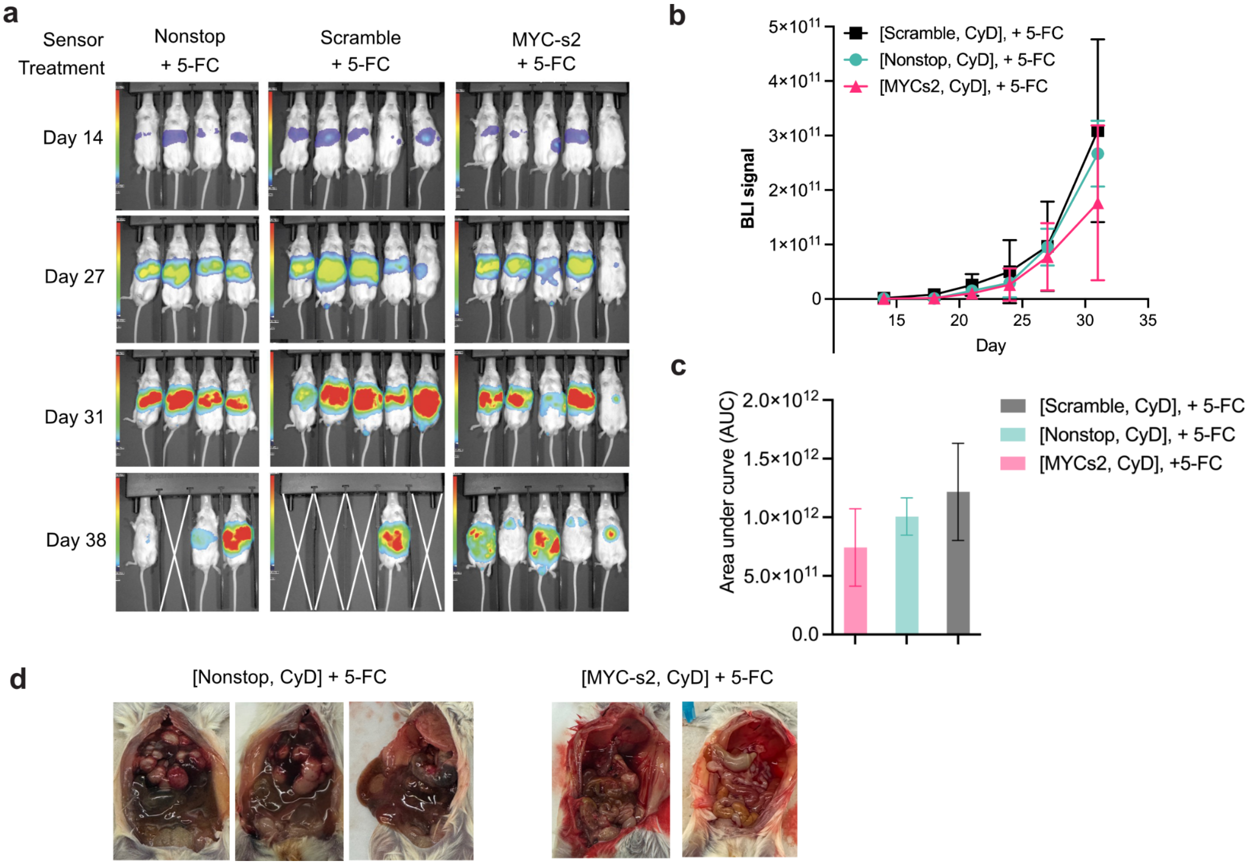


**Supplementary Figure 8. CRC liver metastasis model with AAV-delivered ORCA. a**, Bioluminescence imaging of mice in three groups on different days. **b**, Quantification of bioluminescence signals over time. **c**, Area under the curve (AUC) of bioluminescence signals in the indicated groups in **b**. **d**, Representative images of dissected mice treated with AAV1 encoding RADAR-CyD with nonstop or MYC-s2 sensors. N = 4 mice in the Nonstop group and N = 5 mice in the MYC-s2 and Scramble groups. Error bars represent standard deviation in **b** and **c**.

### Supplementary Table 1. Information of identified RADAR sensors

| Sensor | Length (nt) | Target region | Sequence (5' to 3') | GC content | RNAFold MFE (kcal/mol) | Source |
|---|---|---|---|---|---|---|
| MYC-s1 | 180 | 3'UTR | ATGTACTTAAATAAAAAAAATcAGG<br>GTTTAcAGTACCTAcAATACcAGGT<br>ACTATATACTAGGATTGAAATTCT<br>GTGTAACTGCTATAAACGTTTTAT<br>TAAAGTTATTTACATTcAATaGCAA<br>TATTTACAGAGAAACATTGTGcAA<br>ATCTTAAAATTTTTTAAAAACAATT<br>CTcAAATA | 24.4% | -27.2 | this study |
| MYC-s2 | 180 | 3'UTR | GGAAAACTCTTGCCTTTCTTAAGA<br>CCACCCTCTGAAGGGGCAATTGA<br>TGAAAACAAACAGGGATGGTGGT<br>GGTTGGGAGGGGGAAGAAACGAA<br>ACAAAACATTCACAACTTAAGATT<br>TaGCTCAATGATATATTTGCCAGT<br>TATTTTATTTTTTCTAAAAACAATA<br>GAAAAAAATCAACT | 35.6% | -37.0 | this study |
| FGFR2-s1 | 183 | 3'UTR | GTGCTCTGTAAGTGTGTGCTGAC<br>ATcAATCTTCTCCAATTATTACAAA<br>ATTAACAAGGAAGGCAGAACGCA<br>CGTCCACCTTGAGTCCTACTaGTC<br>CACAGCCAGTACGCACGGCAGGT<br>cAGAGGGGTTACATGGTGGCTTG<br>TGGCAGTCCACTGCTCCAGAAAC<br>CTTCTTCTCCTCCTGGGGA | 50.8% | -49.4 | this study |
| KRAS-s1 | 93 | 3'UTR | TTTTCACAGGCATTGCTAGTTCAA<br>AAACCAAAACTCTGGGAATACTaG<br>CACTTAGAGGAAAAAAAAACTTCC<br>ACTGTCATTTTAAAATcAGCA | 34.4% | -15.0 | this study |
| KRAS-s2 | 183 | 3'UTR | ATGCACCAAAAGCCCCAAGACAG<br>AAATCTTAGGTATTCAGTTTCTTTT<br>TCACAGGCATTGCTAGTTCAAAAA<br>CCAAAACTCTGGGAATACTaGCAC<br>TTAGAGGAAAAAAAAACTTCCACT<br>GTCATTTTAAAATcAGCATTTAAGG<br>TAAAAGCTAACAGTCTGCATGGAG<br>CAGGAAAAAAATTA | 35.5% | -30.4 | this study |
| CDX2-s1 | 93 | 3'UTR | AAAAATAGTTGCAGACAAAATCTG<br>AAGATAAATACAGAATCTGAAATC<br>TaGAAAGCTCATTTATCTCTTTTTC<br>TTTTTAGAAAAAATTCAACT | 24.70% | -14.2 | this study |
| PCNA | 219 | Exon 1 | ctggtgaggttcacgcccatggccaggttgcggt<br>cgcagcgggaggtgtcgaagccctcagaccgc<br>agggtgagctgcaccaaagagacgtaggacga<br>gtccatgctctgcaggtttacaccgctggagctaa<br>tatcccagcaggcctcgttgatgaggtccttgagt<br>gcctccaacaccttcttgaggatggagccctgga<br>ccaggcgcgcctcgaac | 61.20% | -87.4 | CellREADR |

**Supplementary Table 2. Primers used in this work**

| **qPCR primers** | | |
|---|---|---|
| **Target** | **Forward** | **Reverse** |
| ACTB | CGTCCACCGCAAATGCTT | GTTTTCTGCGCAAGTTAGGTTTTGT |
| c-MYC | GTCCTGAGCAATCACCTATGAA | AAGTCCAATTTGAGGCAGTTTAC |
| FGFR2 | AGCTACACTGAGCAGGGAGA | AGGGGTTACATGGTGGCTTG |
| KRAS | CCACGGTCATCCAGTGTTGT | CAGGACCACCACAGAGTGAG |
| CDX2 | acagtcgctacatcaccatccg | cctctcctttgctctgcggttc |
| PCNA | CAAGTAATGTCGATAAAGAGGAGG | GTGTCACCGTTGAAGAGAGTGG |
| ADAR1 | TCCGTCTCCTGTCCAAAGAAGG | TTCTTGCTGGGAGCACTCACAC |

### Supplementary Table 3. Plasmids used in this work

| Plasmid number | Plasmid name |
|---|---|
| pMZ008 | sffv-mcherry-sg-cmyc-s2-sg-egfp-3xms2-flp-in |
| pMZ024 | sffv-mcherry-sg-cmyc-s2.1-sg-egfp-3xms2-flp-in |
| pMZ025 | sffv-mcherry-sg-cmyc-s2.2-sg-egfp-3xms2-flp-in |
| pMZ065 | sffv-mcherry-GSGP2A.cmyc-s1.GSGP2A-egfp-3xms2-flp-in |
| pMZ049 | sffv-mcherry-GSGP2Acmyc-s2GSGP2A-egfp-3xms2-flp-in |
| pMZ061 | sffv-mcherry-GSGP2A.PCNAs.GSGP2A-egfp-3xms2-flp-in |
| pMZ336 | EF1a-mcherry-GSGP2A-CDX2s1-GSGP2A-egfp-3xms2-flp-in |
| pMZ264 | lenti-EF1a-mcherry-MYCs2-GSGP2A-hCD19_SV40-PuroR |
| pMZ265 | lenti-EF1a-mcherry-scramble.GSGP2A-hCD19_SV40-PuroR |
| pMZ266 | lenti-EF1a-mcherry-nonstop.GSGP2A-hCD19_SV40-PuroR |
| pMZ271 | lenti-EF1a-mcherry-MYCs2-GSGP2A-CyD_SV40-PuroR |
| pMZ272 | lenti-EF1a-mcherry-scramble.GSGP2A-CyD_SV40-PuroR |
| pMZ273 | lenti-EF1a-mcherry-nonstop.GSGP2A-CyD_SV40-PuroR |
| pMZ119 | EF1a-mcherry-GSGP2Acmyc-s2GSGP2A-egfp-3xms2-flp-in |
| pMZ120 | EF1a-mcherry-GSGP2Acmyc-scrambleGSGP2A-egfp-3xms2-flp-in |
| pMZ121 | EF1a-mcherry-GSGP2Acmyc-s1GSGP2A-egfp-3xms2-flp-in |
| pMZ132 | EF1a-mcherry-GSGP2A.MYCs2nonstop.2GSGP2A-egfp-3xms2-flp-in |
| pMZ192 | (PB) CAGG-ADAR1p150-SV40-HygroR |
| YH130 | lenti-EF1a-ADAR1-T2A-hygroR |
| YH132 | lenti-EF1a-mcherry-KRAS3'UTR183-s2-GSGP2A-EGFP_SV40-PuroR |
| YH178 | EF1a-mcherry-FGFR23'UTR183-s1-GSGP2A-rtTA-BghplA |
| YH240 | EF1a-mcherry-KRAS3'UTR90-s1-GSGP2A-rtTA-BghplA |
| YH241 | EF1a-mcherry-KRAS23'UTR183-s2-GSGP2A-rtTA-BghplA |
| YH243 | EF1a-mcherry-CDX2-s1-MYC-s2-GSGP2A-rtTA-BghplA |
| YH248 | AAV-EF1a-mcherry-MYCs2-GSGP2A-CyD-WPRE |
| YH249 | AAV-EF1a-mcherry-scramble-GSGP2A-CyD-WPRE |
| YH250 | AAV-EF1a-mcherry-nonstop-GSGP2A-CyD-WPRE |
| YH251 | AAV-EF1a-mcherry-MYCs2-GSGP2A-GFP-WPRE |
| YH252 | CMV-TO-BFPmut-CDX2_3UTR (FLP-IN) |
| YH259 | CMV-TO-IVT-cmyc-s2GSGP2A-NanoLuc-3xms2-flp-in |
| YH260 | CMV-TO-IVT-cmyc-scramble-GSGP2A-NanoLuc-3xms2-flp-in |
| YH261 | CMV-TO-IVT-cmyc-nonstop.GSGP2A-NanoLuc-3xms2-flp-in |
| YH262 | T7-AU-VEE-fluc-cmyc-s2GSGP2A-NanoLuc_XTEN |
| YH263 | T7-AU-VEE-fluc-cmyc-scramble-GSGP2A-NanoLuc_XTEN |
| YH264 | T7-AU-VEE-fluc-cmyc-nonstop.GSGP2A-NanoLuc_XTEN |
| YH265 | CMV-TO-IVT-UTR_Nluc-cmyc-s2.GSGP2A-Ffluc |
| YH266 | CMV-TO-IVT-UTR_Nluc-scramble.GSGP2A-Ffluc |
| YH267 | CMV-TO-IVT-UTR_Nluc-nonstop.GSGP2A-Ffluc |
